# Stop the crop: insights into the insecticidal mode of action of cinnamodial against mosquitoes

**DOI:** 10.1101/2020.05.30.125203

**Authors:** Megha Kalsi, Anton Walter, Beenhwa Lee, Andrew DeLaat, Renata Rusconi Trigueros, Katharina Happel, Rose Sepesy, Bao Nguyen, Preston K. Manwill, H. Liva Rakotondraibe, Peter M. Piermarini

## Abstract

Cinnamodial (CDIAL) is a drimane sesquiterpene dialdehyde found in the bark of Malagasy medicinal plants (*Cinnamosma* species; family Canellaceae). We previously demonstrated that CDIAL was insecticidal, antifeedant, and repellent against *Aedes aegypti* mosquitoes. The goal of the present study was to generate insights into the insecticidal mode of action for CDIAL, which is presently unknown. We evaluated the effects of CDIAL *in vitro* on the contractility of the ventral diverticulum (crop) in adult female *Ae. aegypti*. The crop is a food storage organ surrounded by visceral muscle that spontaneously contracts *in vitro*. We found that CDIAL completely inhibited spontaneous contractions of the crop as well as those stimulated by the agonist 5-hydroxytryptamine. Several derivatives of CDIAL with known insecticidal activity also inhibited crop contractions. Morphometric analyses of crops suggested that CDIAL induced a tetanic paralysis that was dependent on extracellular Ca^2+^ and inhibited by Gd^3+^, a non-specific blocker of plasma membrane Ca^2+^ channels. Screening of numerous pharmacological agents revealed that a Ca^2+^ ionophore (A23187) was the only compound other than CDIAL to completely inhibit crop contractions via a tetanic paralysis. Taken together, our results suggest that CDIAL inhibits crop contractility by elevating intracellular Ca^2+^ through the activation of plasma membrane Ca^2+^ channels thereby leading to a tetanic paralysis, which may explain the insecticidal effects of CDIAL against mosquitoes. Our pharmacological screening efforts also revealed the presence of two regulatory pathways in mosquito crop contractility not previously described: an inhibitory glutamatergic pathway and a stimulatory octopaminergic pathway. The latter was also completely inhibited by CDIAL.

## Introduction

Vector control with chemical insecticides is common strategy employed in the management of mosquito-borne diseases. Unfortunately, the overuse of chemicals with limited and sometimes redundant modes of action has strongly selected for insecticide resistance in mosquito populations. Thus, to maintain a useful chemical arsenal for controlling mosquitoes and their transmission of pathogens, the discovery and development of insecticides with novel modes of action are required (Hemingway et al., 2006; Shaw and Catteruccia, 2019). Plant secondary metabolites are valuable resources for discovering and developing new insecticides (Lorsbach et al., 2019; Norris et al., 2018; Sparks et al., 2019). Notably, several successful classes of insecticides exploit similar modes of action as plant secondary metabolites, including pyrethroids, neonicotinoids, carbamates, and diamides.

Cinnamodial (CDIAL) is a drimane-type, dialdehyde-bearing sesquiterpene isolated from the bark of Malagasy medicinal plants (*Cinnamosma* species of the family Canellaceae). We recently demonstrated that CDIAL was capable of killing larval and adult female mosquitoes (*Aedes aegypti, Anopheles gambiae, Culex pipiens*) (Inocente et al., 2018). Notably, the insecticidal potency of CDIAL was similar against pyrethroid-susceptible and pyrethroid-resistant lines of *Ae. aegypti*, suggesting a unique mode of action compared to pyrethroids (Inocente et al., 2018). CDIAL was also antifeedant and repellent against adult female *Ae. aegypti* and an agonist of mosquito transient receptor potential ankyrin 1 (TRPA1) channels (Inocente et al., 2018). Although agonism of TRPA1 channels by CDIAL was responsible for its antifeedant activity, and likely contributed to the repellent effects against mosquitoes, TRPA1 modulation was not required for the insecticidal activity of CDIAL (Inocente et al., 2018). Consistent with this notion, the insecticidal and antifeedant activities of various natural and semi-synthetic derivates of CDIAL often do not correlate (Inocente et al., 2019; Manwill et al., 2020). Thus, the mode of insecticidal activity for CDIAL is currently unknown and appears to be independent from its mode of antifeedant activity, which is by TRPA1 activation.

The goal of the present study was to generate insights into the insecticidal mode of action for CDIAL by assessing the effects it has on the contractile activity of the ventral diverticulum (crop) in adult female *Ae. aegypti*. The crop is a diverticulation of the foregut in adult flies (order Diptera) and plays a key role in food storage (Stoffolano and Haselton, 2013). The lumen of the crop consists of a cuticular-lined epithelium surrounded on the serosal side by a meshwork of visceral muscle; the latter undergoes rhythmic peristaltic contractions when the crop is full (Stoffolano and Haselton, 2013). The mosquito crop is the primary receptacle of imbibed sugar solutions, such as floral nectars in the field or sucrose in the laboratory (Day, 1954; Trembley, 1952). Peristaltic contractions of the crop empty its contents into the midgut for digestion and absorption. We demonstrated previously that the crop of *Ae. aegypti* undergoes spontaneous peristaltic contractions *in vitro* that are Ca^2+^-dependent, stimulated by 5-hydroxytryptamine (5-HT), and inhibited by a chemical mimic of myosupressins, benzethonium chloride (Calkins et al., 2017). Here we demonstrate that CDIAL completely inhibits the spontaneous and 5-HT stimulated contractions of the crop *in vitro*. In addition, by comparing the inhibitory effects of CDIAL with other pharmacological agents, a hypothesis is generated as to the mode of insecticidal action for CDIAL against mosquitoes. Furthermore, this study provides new insights into the physiological regulation of crop contractility in mosquitoes.

## Materials and Methods

### Mosquitoes

Eggs of *Ae. aegypti* were obtained originally from the Malaria Research and Reference Reagent Resource Center (MR4) as part of the BEI Resources Repository (Liverpool strain; LVP-IB12 F19, deposited by M.Q. Benedict). Mosquitoes were reared to adults as described previously (Piermarini et al., 2011) and held in environmentally-controlled chambers (28°C, 80% relative humidity, 12 h:12 h light:dark cycle). Tetramin flakes (Melle, Germany) were pulverized and fed to larvae daily. Adults were maintained on 10% sucrose *ad libitum*. To generate additional eggs, adult females were periodically fed defibrinated rabbit blood (Hemostat Laboratories, Dixon, CA) with a membrane feeder (Hemotek, Blackburn, UK).

### Isolation of mosquito crops and hindguts

Crops were isolated from non-blood fed adult females (3-10 days post-emergence) under mosquito Ringer’s solution as previously described (Calkins et al., 2017). If a crop was leaking its contents, filled with air bubbles, or not contracting spontaneously, then it was discarded. The Ringer’s solution consisted of the following in mM: 150 NaCl, 3.4 KCl, 1.7 CaCl2, 1.8 NaHCO3, 1 MgCl2, 5 Glucose, and 25 HEPES. The pH of the Ringer’s solution was adjusted to 7.1 with 1 N NaOH and its osmolality verified to be 330 ± 5 mOsmol/kg with a vapor pressure osmometer (Wescor, Logan, UT). In some experiments we used a ‘zero Ca^2+^’ Ringer’s solution in which CaCl2 was excluded and 1 mM EGTA (Thermo Fisher Scientific, Waltham, MA) was added.

The hindguts from adult females were also utilized for some experiments. In brief, the alimentary canal was isolated under Ringer’s solution by tugging the last abdominal segment with fine forceps (Dumont #5; Fine Science Tools, Inc., Foster City, CA). The ileum of the hindgut was isolated by pinching it off from its junctions with the midgut and rectum with fine forceps.

### Chemicals

All natural drimane sesquiterpenes were isolated from medicinal plants and their structures were determined as previously described (He et al., 2017; Inocente et al., 2019; Inocente et al., 2018; Manwill et al., 2020). In brief, cinnamodial (CDIAL), polygodial (POLYG), ugandensolide (UGAN), cinnamosmolide (CMOS), cinnamolide (CML), drimenin (DRIM), and capsicodendrin (CPCD) were isolated from the bark of *Cinnamosma fragrans* Baill (Canellaceae), whereas warburganal (WARB) was isolated from the bark of *Warburgia ugandensis* Sprague (Canellaceae) (Inocente et al., 2019). Their structures are shown in Supplemental Fig. 1. Semi-synthetic derivatives of CDIAL were produced as described previously (Manwill et al., 2020). Herein, the derivatives are referred to as CD02, CD03, CD04, CD05, CD06, CD08, CD09, CD10, CD11, CD12. Their structures are shown in Supplemental Fig. 2. All drimane sesquiterpenes and CDIAL derivatives were dissolved in dimethyl sulfoxide (DMSO, Thermo Fisher Scientific) at 100-times the final assay concentrations. In some experiments CPCD was dissolved in acetone (Thermo Fisher Scientific) at 100-times the final assay concentration. The other pharmacological agents utilized in the present study and their respective solvents are listed in Supplemental Table 1.

### *Measurements of crop and hindgut contraction rates* in vitro

We utilized an *in vitro* assay previously established for mosquito crops (Calkins et al., 2017). In brief, an isolated crop or hindgut was transferred to a single well of a 96-well microtiter plate (USA Scientific, Ocala, FL) containing 100 μl of Ringer’s solution where it rested for 5 min. After the rest period, the frequency of crop or hindgut contractions was counted by eye under a stereoscope every other minute for 5 min (i.e., during the first, third, and fifth minute following the rest period). These contraction frequencies were averaged together and considered the ‘control’ rate. After the final control observation, 1 μl of Ringer’s solution was removed from the well and replaced with 1 μl of a treatment solution (see *Chemicals*), followed by gentle mixing via pipetting. After a 2 min incubation period, the frequency of contractions was counted again every other minute for 5 min. These contraction frequencies were averaged together and referred to as the ‘treatment’ rate. Our previous study (Calkins et al., 2017) and preliminary trials in the present study found that the solvents did not affect contraction rates at a 1% final assay concentration. In some experiments a ‘pre-treatment’ period with CDIAL, 5-HT, or octopamine (OA) preceded the ‘treatment’ period.

### Crop morphometric analyses

To determine if treatments affected crop shape, crops were prepared for the *in vitro* assay as described above and recorded under a stereoscope (40x total magnification) with a digital camera (Dino-Lite, Torrance, CA). The recordings consisted of a 5 min control period followed by a 5 min treatment period. Video recordings were played back on a personal computer using the ‘Movies & TV’ app in *Windows 10* (Microsoft, Redmond, WA). Three still images were saved as JPG files during the control period when the crop was in a fully relaxed/flaccid state. Another still image was saved five minutes into the treatment period.

For each crop, the JPG files were opened with *Fiji* image analysis software (Schindelin et al., 2012). The perimeter of each crop was traced carefully using the polygon selection tool and the edges were rounded using the ‘Fit Spline’ function. The following steps under the ‘Edit’ menu were then performed to simplify the image: ‘Clear Outside’; ‘Clear’; and ‘Fill’. The simplified image was converted to ‘8-bit’ format to make it compatible with the *MorphoLibJ* morphometrics plug-in (Legland et al., 2016), which was used to fit an object-oriented bounding box (OOBB) to the crop. The length of the OOBB was a measure of the crop’s distal to proximal length; the width of the OOBB was a measure of its girth. An ‘oriented box elongation ratio’ (OBER) was calculated by dividing the width of the OOBB by its length. A ratio of 1.0 indicated a circular shape; a ratio <1.0 indicated a more elongate shape (Wirth, 2004). The three OBER measurements from the control period were averaged together.

### Statistics

All statistical analyses were performed in *Prism* 6.07 (GraphPad Software Inc., La Jolla, CA). The specific tests utilized are described in the text of the **Results** and figure legends.

## Results

### CDIAL inhibits the frequency of crop contractions

We first determined the effects of CDIAL on the frequency of spontaneous crop contractions. Initial experiments found that adding 0.1 μM CDIAL to the bath reduced the spontaneous contraction rates by ~15% (Fig. 1A). However, a 10-fold higher concentration of CDIAL (1 μM) did not affect the frequency of crop contractions (Fig. 1B). On the other hand, a 100-fold higher concentration (10 μM CDIAL) consistently and completely inhibited the spontaneous contractions (Fig. 1C).

**Fig. 1.**
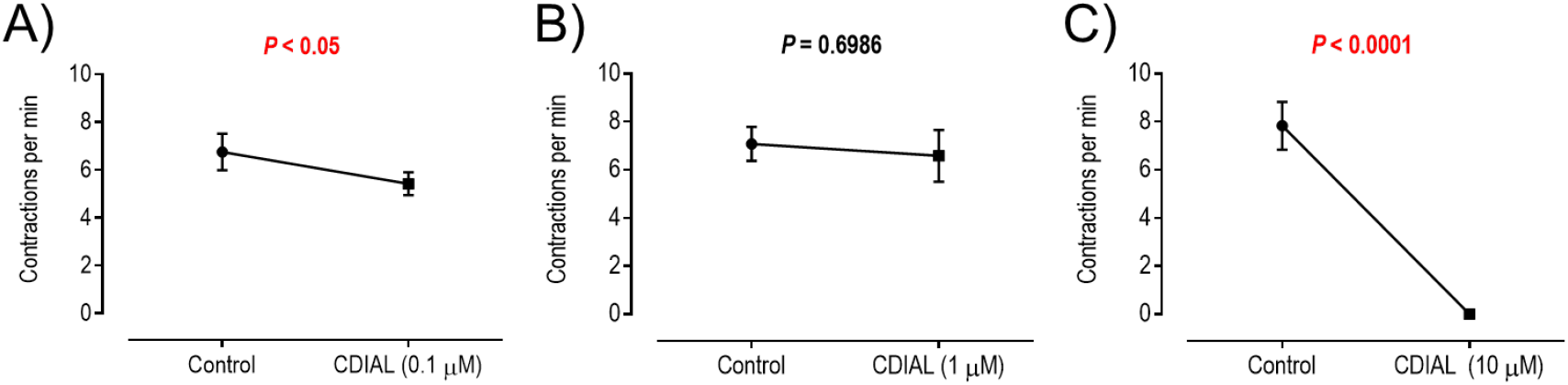
Effects of CDIAL on spontaneous contractions of the crop. A) 0.1 μM (N = 11), B) 1 μM (N = 12), C) 10 μM (N = 10). Values are means ± SEM. *P* values from paired t-tests are indicated with significant values in red text.

We next determined whether 10 μM CDIAL also inhibited crops stimulated with 5-hydroxytryptamine (5-HT). We previously demonstrated that 5-HT is a potent agonist of mosquito crop contractions (Calkins et al., 2017). Here we confirm that 100 nM 5-HT increased the frequency of crop contractions (Fig. 2A). Subsequent treatment with 10 μM CDIAL completely inhibited crop contractions (Fig. 2A). In contrast, if 1% DMSO (solvent for CDIAL) was added to the bath, then the 5-HT-stimulated contractions continued unperturbed (Supplemental Fig. 3A). We also tested whether CDIAL treatment prevented the stimulatory effects of 5-HT. As shown in Fig. 2B, if crops were first treated with 10 μM CDIAL, which completely stopped the spontaneous contractions, then subsequent addition of 100 nM 5-HT had no effect (Fig. 2B). We also tested CDIAL on the adult female hindgut to determine if it inhibited spontaneous contractions of another visceral muscle. As shown in Supplemental Fig. 3B, 10 μM CDIAL inhibited the spontaneous contraction rates of the hindgut by ~92%.

**Fig. 2.**
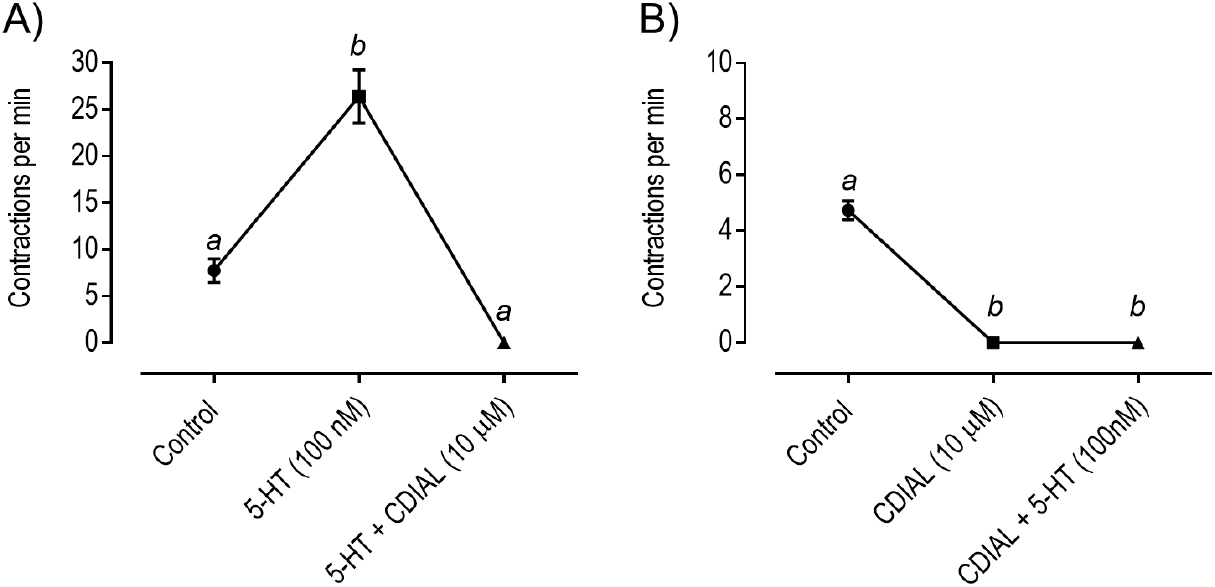
Effects of CDIAL on 5-HT-stimulated peristaltic contractions of the crop. A) Crops were stimulated with 100 nM 5-HT before addition of 10 μM CDIAL (N = 5). B) Crops were treated with 10 μM CDIAL before addition of 100 nM 5-HT (N = 5). Values are means ± SEM. Lower-case letters indicate statistical categorization of the means as determined by a repeated measures one-way ANOVA and Tukey’s multiple comparisons test (P < 0.05).

We next investigated whether several natural or semi-synthetic derivatives of CDIAL, which we have previously assessed for insecticidal and antifeedant activities against mosquitoes (Inocente et al., 2019; Inocente et al., 2018; Manwill et al., 2020), affected the frequency of crop contractions. First, we confirmed that 10 μM CDIAL completely inhibited the spontaneous contractions (Supplemental Fig. 4A; Fig. 3). Among the natural monomeric derivatives tested (all at 10 μM), the dialdehydes POLYG and WARB inhibited crop contraction rates by ~82 and 65%, respectively (Supplemental Fig. 4B-C; Fig. 3), whereas the lactone derivative UGAN stimulated crop contraction rates by ~50% (Supplemental Fig. 4D; Fig. 3). The other lactone derivatives, CMOS, CML, and DRIM, did not affect the frequency of crop contractions (Supplemental Fig. 4E-G). We also tested CPCD, a natural dimeric derivative of CDIAL with only one aldehyde group (Supplemental Fig. 1). CPCD completely stopped crop contractions when it was dissolved in DMSO (Supplemental Fig. 4H; Fig. 3). However, when it was dissolved in acetone, CPCD did not affect contraction rates (Supplemental Fig. 4I). Among the semi-synthetic CDIAL derivatives tested (all at 10 μM), CD02, CD03, CD04, CD05, CD06, and CD10 showed no detectable effects (Supplemental Fig. 4A-E, H). On the other hand, CD08, CD09, CD11, and CD12 inhibited crop contraction rates by ~83, 58, 20, and 22%, respectively (Supplemental Fig. 5F-G, I-J; Fig. 3).

**Fig. 3.**
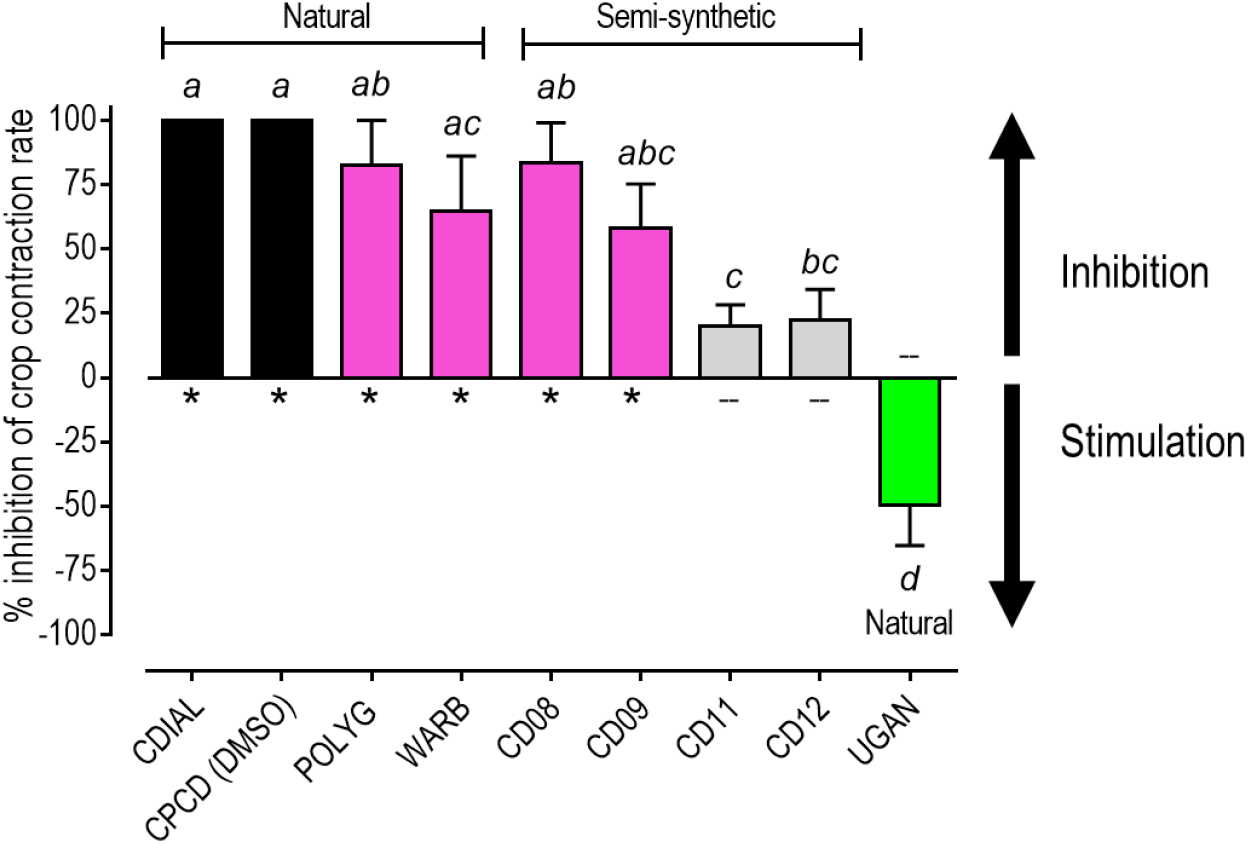
Percent inhibition of spontaneous crop contraction rates by CDIAL derivatives with significant effects. Values are means ± SEM, based on the percent inhibition of control contraction rates in Supplemental Figs. 4&5. Lower-case letters indicate statistical categorization of the means as determined by a repeated measures one-way ANOVA and Tukey’s multiple comparisons test (P < 0.05). Colors indicate qualitative categorization of effects: black = 100% inhibition; magenta = moderate-strong inhibition (>50%); gray = weak inhibition (<50%); and green = stimulation. ‘*’ or ‘--’ respectively indicate compounds with or without previously demonstrated insecticidal activity against *Ae. aegypti* (Inocente et al., 2019; Inocente et al., 2018; Manwill et al., 2020).

In summary, among compounds that affected crop contraction rates, CDIAL and CPCD (dissolved in DMSO) were the most efficacious inhibitors (Fig. 3). POLYG, WARB, CD08, and CD09 were moderate inhibitors causing ~50-82% inhibition. CD11 and CD12 were weak inhibitors causing <25% inhibition (Fig. 3). On the other hand, UGAN was an activator (Fig. 3). All of the moderate-strong or complete inhibitors of crop contractions rates were previously shown to be insecticidal (‘*’ in Fig. 3) against adult female and/or larval *Ae. aegypti* (Inocente et al., 2019; Inocente et al., 2018; Manwill et al., 2020).

### Modulators of neurotransmitter receptors, but not acetylcholinesterase, affect crop contraction rates

Previous studies have found that plant-derived insecticides modulate neurotransmitter receptors, such as Cys-loop gated ion channels and G protein-coupled receptors, and/or inhibit acetylcholinesterase (Anderson and Coats, 2012; Furutani et al., 2014; Liao et al., 2016; Tong et al., 2013). Thus, we tested whether the inhibition of crop contraction frequency by CDIAL might be explained by modulation of similar mechanisms. First, we screened the effects of various neurotransmitters (all at 1 mM) on crop contraction rates. As shown in Supplemental Fig. 6A-C and Fig. 4, glutamate, acetylcholine, and GABA inhibited the frequency of crop contractions by ~80, 30, and 10%, respectively. On the other hand, OA increased the frequency of crop contractions by ~250% (Supplemental Fig. 6D; Fig. 4). Neither histamine nor aspartate affected crop contraction rates (Supplemental Fig. 6E-F). The percent inhibition by glutamate was greater than that of acetylcholine and GABA (Fig. 4).

**Fig. 4.**
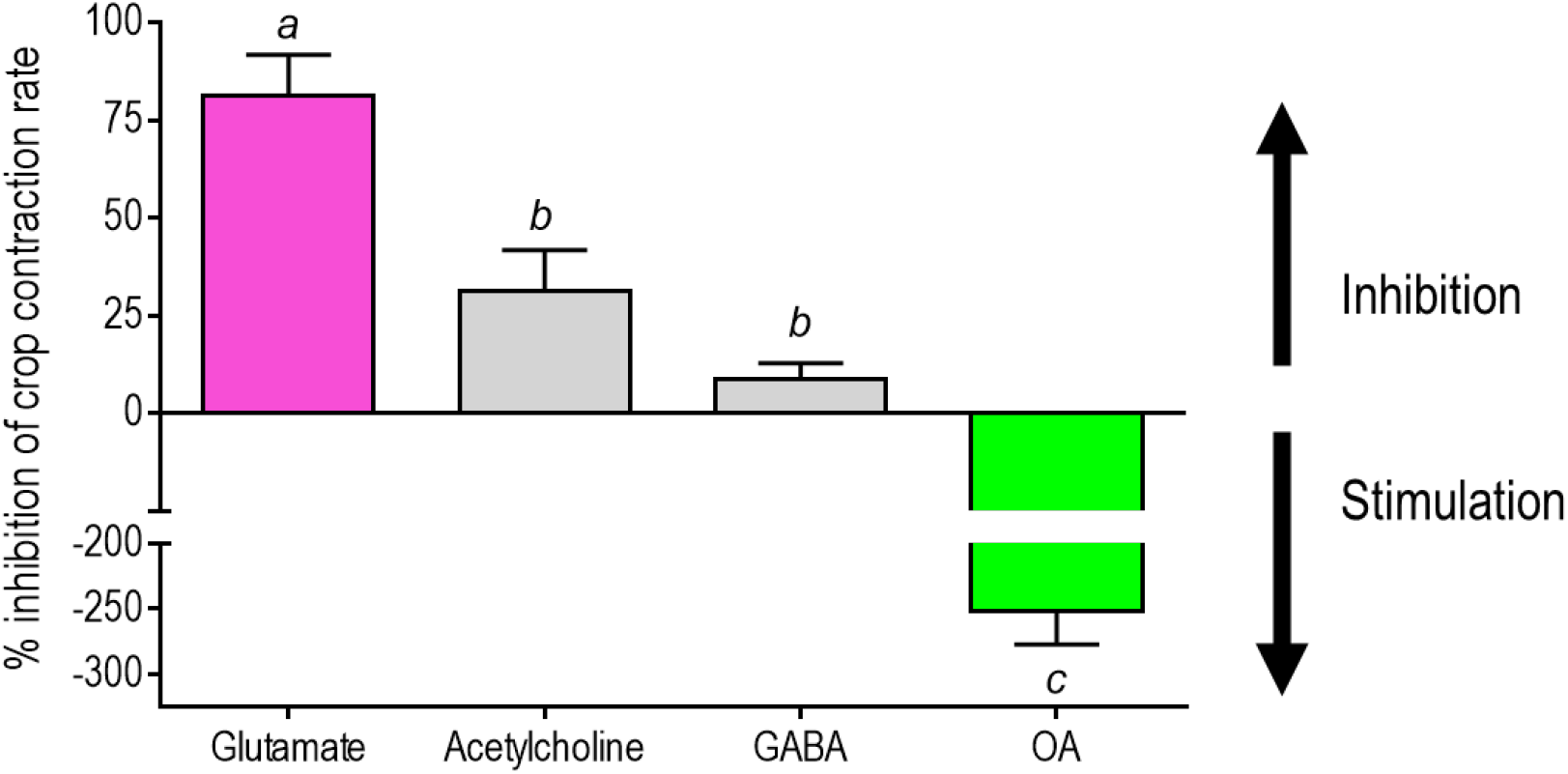
Percent inhibition of spontaneous crop contraction rates by neurotransmitters (1 mM) with significant effects. Values are means ± SEM, based on the percent inhibition of control contraction rates in Supplemental Fig. 6. Lower-case letters and colors respectively indicate statistical and qualitative categorizations of the effects as in Fig. 3.

To further characterize the inhibition of crop contractions by glutamate we screened more discriminating modulators of glutamate receptors. Notably, two agonists of glutamate-gated chloride (GluCl) channels, ivermectin (5 μM) and ibotenate (1 mM), each completely inhibited crop contractions (Supplemental Fig. 7A-B; Fig. 5). Among agonists of ionotropic glutamate receptors tested (all at 1 mM), NMDA inhibited crop contraction rates ~28% (Supplemental Fig. 7C; Fig. 5), whereas neither quisqualate nor kainate affected crop contraction rates (Supplemental Fig. 7D-E).

**Fig. 5.**
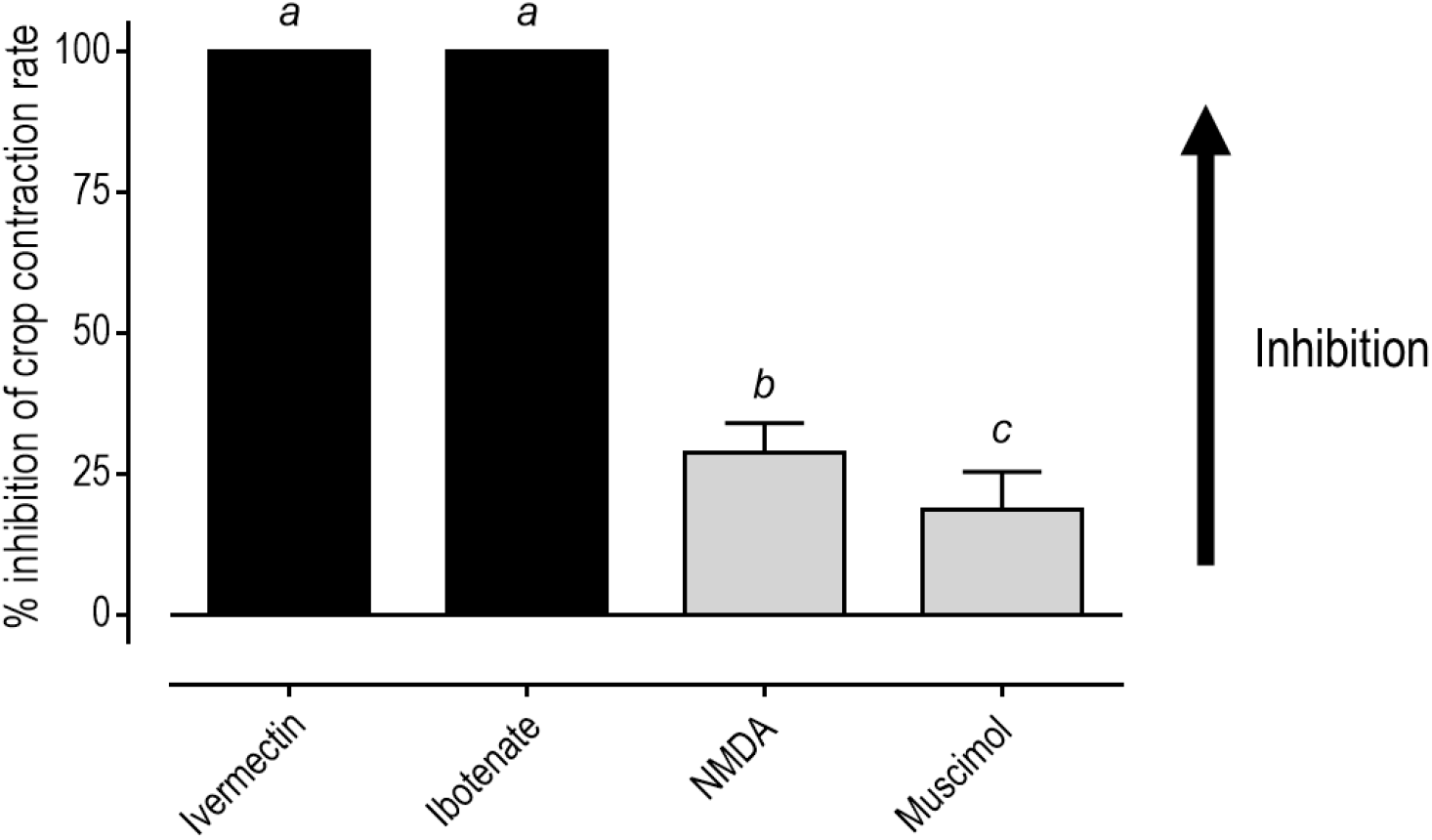
Percent inhibition of spontaneous crop contraction rates by glutamate or GABA receptor agonists with significant effects. All were tested at 1 mM except for ivermectin (5 μM). Values are means ± SEM, based on the percent inhibition of control contraction rates in Supplemental Fig. 7. Lower-case letters and colors respectively indicate statistical and qualitative categorizations of the effects as in Fig. 3.

Given that GABA and acetylcholine inhibited crop contractions (Supplemental Fig. 6B-C; Fig. 4) we tested modulators of GABA and acetylcholine receptors, as well as acetylcholinesterase. As shown in Supplemental Fig. 7F and Fig. 5, muscimol (1 mM), a GABA-receptor agonist, inhibited crop contractions by ~20%, which was similar in magnitude to 1 mM GABA (Fig. 4). On the other hand, pilocarpine (10 μM) and imidacloprid (10 μM), agonists of muscarinic and nicotinic acetylcholine receptors, respectively, did not affect the frequency of crop contractions (Supplemental Fig. 8A-B). Acetylcholinesterase inhibitors (100 μM carbaryl, 10 μM propoxur, 10 μM carvacrol) also did not affect crop contraction rates (Supplemental Fig. 8C-E).

Given the stimulatory effects of 1 mM OA (Supplemental Fig.6D; Fig. 4), we tested amitraz, an agonist of insect OA receptors. As shown in Fig. 6A, 10 μM amitraz increased the frequency of crop contractions. We next determined whether OA was stimulatory at a lower concentration (10 μM) and if CDIAL would inhibit the OA-stimulated crop contraction rates. As shown in Fig. 6B, 10 μM OA stimulated crop contraction frequency and subsequent addition of 10 μM CDIAL completely inhibited crop contractions. On the other hand, if crops were treated with 1% DMSO (CDIAL solvent) instead of CDIAL, then the OA-stimulated contraction frequency was unaffected (Supplemental Fig. 9). If crops were first treated with 10 μM CDIAL, which stopped the spontaneous contractions, subsequent addition of OA had no effect (Fig. 6C).

**Fig. 6.**
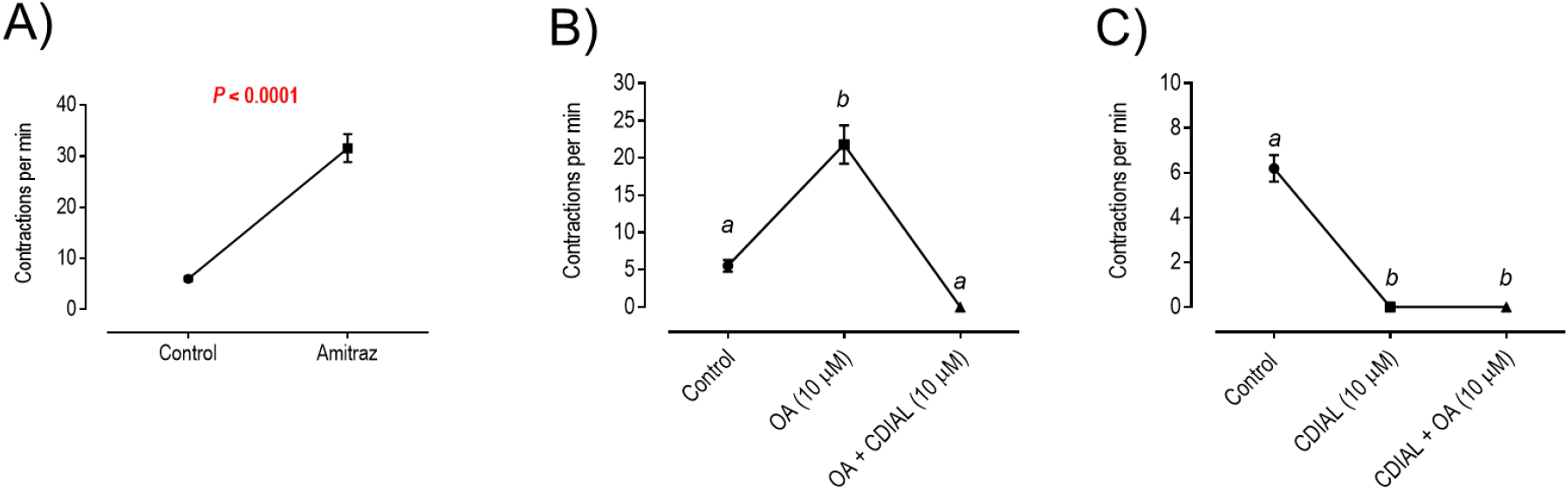
Validation of an octopaminergic stimulatory pathway and its inhibition by CDIAL in the crop. A) Effects of 10 μM amitraz on spontaneous peristaltic contractions of the crop. Values are means ± SEM. *P* value from paired t-test is indicated. N = 9. B) Crops were stimulated with 10 μM OA before addition of 10 μM CDIAL (N = 5). C) Crops were treated with 10 μM CDIAL before addition of 10 μM OA (N = 5). In panels B & C, values are means ± SEM, and lower-case letters indicate statistical categorization of the means as determined by a repeated measures one-way ANOVA and Tukey’s multiple comparisons test (P < 0.05).

### Modulators of plasma membrane Ca^2+^ and TRP channels affect crop contraction frequency

We next tested whether the inhibitory effects of CDIAL on crop contractions could potentially be explained by modulating plasma membrane Ca^2+^ channels. Previously we have shown that extracellular Ca^2+^ is required for the spontaneous contractions of the mosquito crop (Calkins et al., 2017), suggesting the involvement of plasma membrane Ca^2+^ channels in myogenic activity. In the present study, we tested the effects of generic blockers of Ca^2+^-channels: Gd^3+^, SKF-96365, or nifedipine at 100 μM. Each blocker completely inhibited the spontaneous crop contractions (Supplemental Fig. 10A-C; Fig. 7). Additional experiments at a lower concentration (10 μM) revealed complete inhibition of crop contractions by Gd^3+^ (Supplemental Fig. 10D; Fig. 7). Neither 10 μM SKF-96365 nor 10 μM nifedipine affected crop contractions, but the *P* value for nifedipine was near the threshold of significance (Supplemental Fig. 10E-F).

**Fig. 7.**
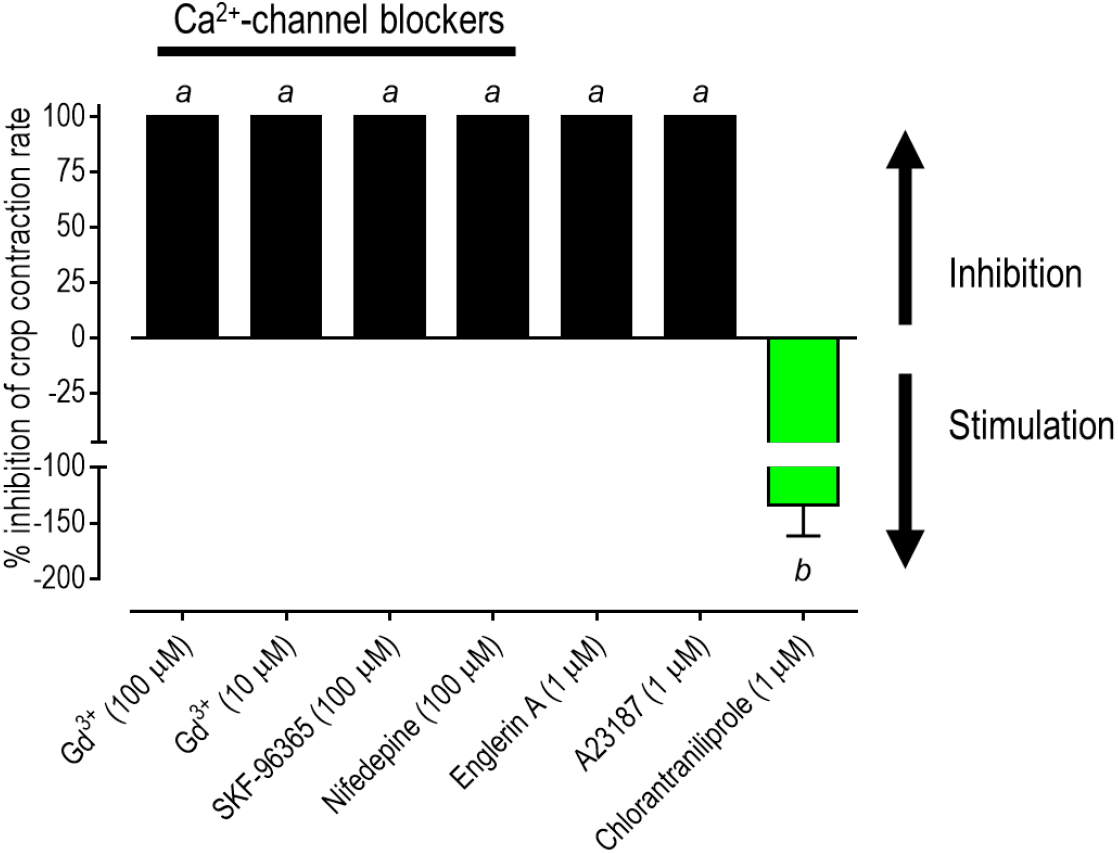
Percent inhibition of spontaneous crop contraction rates by plasma membrane Ca^2+^ channel blockers, a TRP channel agonists (englerin A), a Ca^2+^ ionophore (A23187), or ryanodine receptor agonists (chlorantraniliprole) with significant effects. Values are means ± SEM, based on the percent inhibition of control contraction rates in Supplemental Figs. 10-12. Lower-case letters and colors respectively indicate statistical and qualitative categorizations of the effects as in Fig. 3.

We also tested a few pharmacological agonists of TRP channels, which can mediate the influx of Ca^2+^ and/or other cations when activated. Pymetrozine (500 μM), an agonist of insect TRPV1 channels, did not affect the spontaneous contraction rates (Supplemental Fig. 11A). On the other hand, englerin A (1 μM), a natural product agonist of mammalian TRPC channels (Akbulut et al., 2015; Carson et al., 2015), completely inhibited crop contractions (Supplemental Fig. 11B; Fig. 7).

### Modulators of intracellular Ca^2+^ homeostasis affect crop contraction frequency

We next tested modulators of intracellular Ca^2+^ homeostasis on the spontaneous contractions of the crop. The Ca^2+^ ionophore A23187 (1 μM) completely inhibited the frequency of crop contractions (Supplemental Fig. 12A; Fig. 7). However, neither thapsigargin (1 μM), an inhibitor of intracellular Ca^2+^-ATPases, nor ryanodine (10 μM), an inhibitor of intracellular Ca^2+^ release channels (ryanodine receptors), had significant effects on crop contraction rates (Supplemental Fig. 12B-C). Intriguingly, chlorantraniliprole (1 μM), an agonist of insect ryanodine receptors, increased crop contraction frequency by over 100% (Supplemental Fig. 12D; Fig. 7).

### CDIAL affects the shape of the crop in a Ca^2+^-dependent manner

To obtain additional insights into the mechanisms by which CDIAL may inhibit the spontaneous contractions of the crop we performed morphometric measurements before and after CDIAL treatment. In preliminary observations, we noticed that within 5 min after CDIAL treatment, the longitudinal diameter of the crop (distal to proximal length) contracted, resulting in a change of the crop’s appearance from elongate to circular (Fig. 8).

**Fig. 8.**
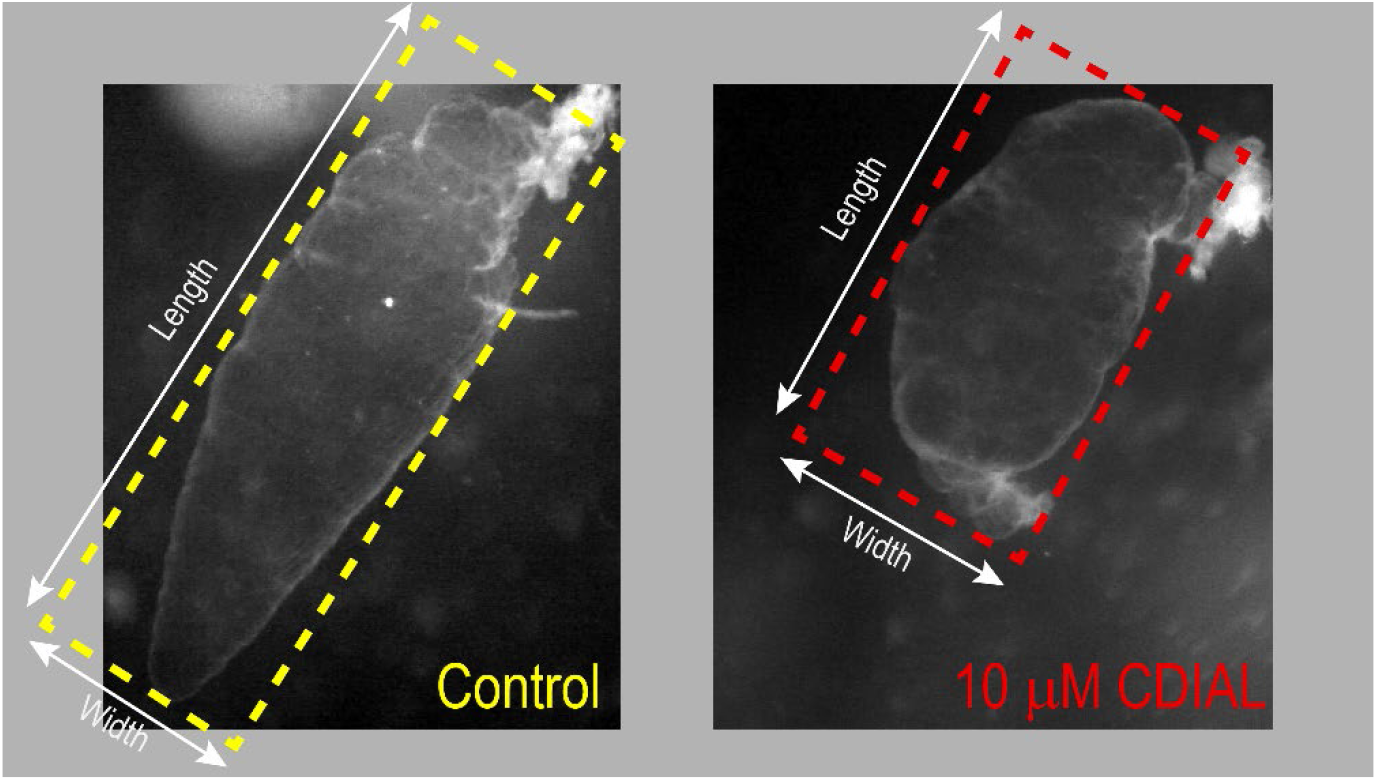
Effects of CDIAL on crop shape. Representative images before (control) and 5 min after treatment of crops with 10 μM CDIAL. Note the contraction in the length of the crop after CDIAL treatment, resulting in a more circular shape. The dashed boxes represent ‘object oriented bounding boxes’ drawn by the ‘MorphoLibJ’ plugin of *Fiji* image analysis software. The ‘oriented box elongation ratio’ is calculated by dividing the width of the box by its length. A ratio of 1.0 indicates a circular shape; a ratio <1.0 indicates a more elongate shape.

This phenomenon was documented and quantified by capturing digital images of crops in their relaxed state before (control) and 5 min after treatment with 10 μM CDIAL. From these images we used digital image analysis software to calculate the ‘oriented box elongation ratio’ (OBER), a simple measure of shape (width/length; Fig. 8). An OBER of 1.0 is indicative of a circular shape, whereas an OBER < 1.0 indicates an elongate shape (Wirth, 2004). Remarkably, 10 μM CDIAL increased the mean OBER of crops from ~0.42 to 0.55, an increase of ~30% (Fig. 9A; Fig. 10). In other words, the crops became more circular in shape. Likewise, 100 μM CDIAL increased the mean OBER of crops from ~0.4 to 0.6, an increase of ~60% (Fig. 9B; Fig. 10).

**Fig. 9.**
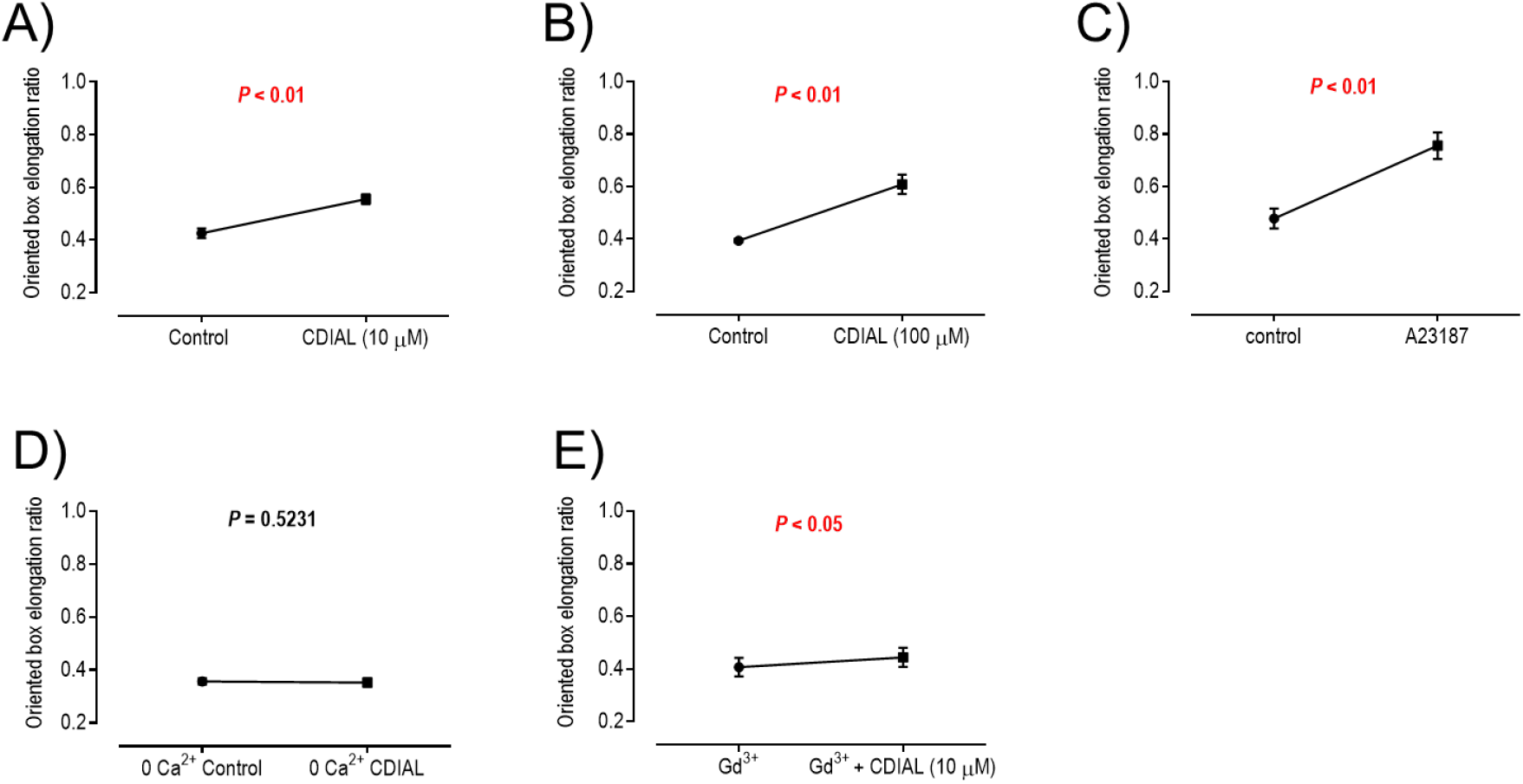
Morphometric effects of CDIAL on the crop. Effects of CDIAL (A, B) or A23187 (C) on the OBER of crops. D) Effects of 10 μM CDIAL on the OBER of crops in a zero Ca^2+^ Ringer’s solution. E) Effects of 10 μM CDIAL on the OBER of crops in the presence of 100 μM Gd^3+^. Values are means ± SEM. *P* values from paired t-tests are indicated. N = 9 for 10 μM CDIAL; N = 10 for 100 μM CDIAL; N =5 for 1 μM A23187; N = 9 for 0 Ca^2+^ experiments; N= 7 for Gd^3+^ experiments.

**Fig. 10.**
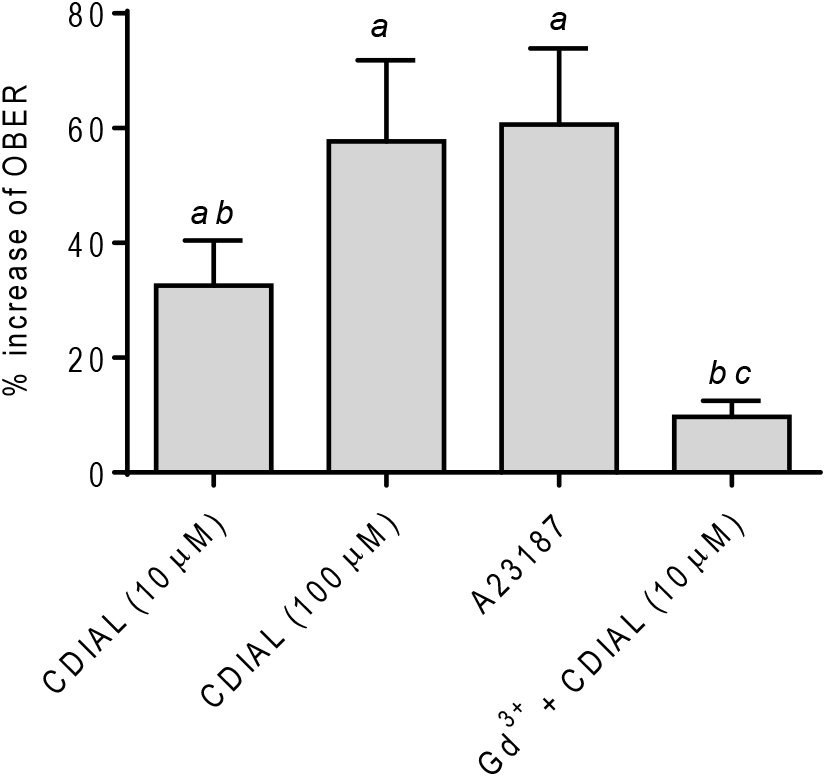
Percent increase of the OBER of crops induced by CDIAL and A23187. Values are means ± SEM based on the percent increase in OBER from the control in Fig. 9. Lower-case letters indicate statistical categorization of the means as determined by a repeated measures one-way ANOVA and Tukey’s multiple comparisons test (P < 0.05).

We next determined whether the pharmacological modulators that completely inhibited crop contractions (Figs. Fig. 4, Fig. 5, & Fig. 7) also affected crop shape like CDIAL. Among these compounds, only A23187 affected the OBER of crops, increasing it by ~60% (Fig. 9C; Fig. 10). The other compounds did not affect the OBER of crops (Supplemental Fig. 13). The similar effects of CDIAL and A23187 suggest that the shape change induced by CDIAL might involve an increase of intracellular Ca^2+^. Thus, we next tested the effects of CDIAL in a Ringer’s solution with nominal Ca^2+^. As shown in Fig. 9D, when a zero Ca^2+^ Ringer’s solution was used, 10 μM CDIAL did not affect the OBER of crops. To determine if plasma membrane Ca^2+^-channels contributed to the effects of CDIAL on crop shape we added 100 μM Gd^3+^ to the Ringer’s solution. In the presence of Gd^3+^, 10 μM CDIAL increased the OBER of crops, but only by ~10% (Fig. 9E; Fig. 10).

## Discussion

To our knowledge, the present study is the first to characterize the physiological effects of drimane sesquiterpenes on an isolated insect tissue. We demonstrated that CDIAL inhibited the spontaneous peristaltic contractions of visceral muscle in the ventral diverticulum (crop) and hindgut of adult female *Ae. aegypti*. In addition, CDIAL completely blocked the enhanced rate of crop contractions stimulated by 5-HT or OA. Thus, CDIAL likely disrupts an essential mechanism mediating the contraction of visceral muscle in the alimentary canal of mosquitoes. It remains to be determined whether CDIAL disrupts the activities of visceral muscle in other mosquito tissues (e.g., ovaries, heart) and/or other muscle types (e.g., flight muscle).

In addition to CDIAL, other natural drimane sesquiterpenes inhibited the spontaneous contractions of the crop, including CPCD, POLYG, and WARB. CPCD is a dimeric form of CDIAL that rapidly converts into CDIAL monomers in the presence of DMSO (Karmahapatra et al., 2018). As such, when CPCD was dissolved in DMSO, it displayed similar inhibition as CDIAL. On the other hand, when CPCD was dissolved in acetone, it lacked effects on crop contractions, which confirms our earlier findings that CPCD has limited bioactivity in its dimeric form (Inocente et al., 2019; Karmahapatra et al., 2018). The strong inhibition of crop contractions by the dialdehydes CDIAL, POLYG and WARB, but not lactone derivatives with similar structures (CMOS, CML, and DRIM), indicate that electrophilic aldehydes are important in the bioactivity of drimane sesquiterpenes against mosquito visceral muscle. We have come to similar conclusions in previous studies regarding the insecticidal, antifeedant, and repellent bioactivities of these molecules (Inocente et al., 2019; Inocente et al., 2018; Manwill et al., 2020). However, in the present study one of the lactone derivatives (UGAN) modestly stimulated contractions of the crop. Moreover, in a previous study, we found that CML possessed comparable antifeedant activity to CDIAL (Inocente et al., 2019). Thus, although aldehydes are generally associated with the bioactivity of drimane sesquiterpenes in mosquitoes, they are not always essential.

In addition to the natural CDIAL derivatives, two semi-synthetic derivatives (CD08, CD09) were prominent inhibitors of crop contractions. CD09 is identical to CDIAL with the exception that the formyl group on C-8 was replaced with a methyl ketone (Supplemental Fig. 1), resulting in a less electrophilic molecule. In previous work we found that CD09 was similarly effective as CDIAL at killing larval and adult female mosquitoes, but a weaker antifeedant, suggesting that insecticidal activity can be maintained if the C-12 aldehyde was replaced with a more stable/less electrophilic moiety (Manwill et al., 2020). The inhibition of crop contractions by CD09 in the present study is consistent with this notion. CD08 is more substantially modified where the C-6 acetoxy group and C-9 aldehyde were each replaced with a ketone carbonyl (hydroquinone) and the C-12 aldehyde was extended with an ethyl formate linker (Supplemental Fig. 1). In previous work we found that CD08 was a superior larvicide compared to CDIAL, but did not kill adult females (Manwill et al., 2020). The inhibition of crop contractions in the present study indicates that CD08 is bioactive against adult female visceral muscle and would be expected to show adulticidal activity. Thus, the results are consistent with our previously suggested notion (Manwill et al., 2020) that the lack of adulticidal activity by CD08 is due to more efficient detoxification and/or excretion in adults compared to larvae. Future studies on the metabolism of CD08 by mosquitoes are necessary to confirm this hypothesis.

Overall, results in the present study along with those in our previous studies characterizing mosquitocidal effects of CDIAL derivatives (Inocente et al., 2019; Inocente et al., 2018; Manwill et al., 2020) suggest there is a general direct correlation between the inhibition of visceral muscle activity in the crop and insecticidal activity of CDIAL derivatives (Fig. 3). In adult mosquitoes, the primary role of the crop is to store sugar meals (e.g., nectar), which are eventually transported to the midgut for digestion and absorption. Emptying of the crop’s contents into the midgut relies on peristaltic contractions mediated by visceral muscle. Thus, adult mosquitoes treated with CDIAL may fail to empty their crops, which may starve mosquitoes of the energy needed for flight and other key physiological functions. Although larval mosquitoes lack a crop, previous studies have demonstrated that visceral muscle of the midgut undergoes 5-HT-dependent peristaltic contractions (Onken et al., 2004), which are likely involved in digestion and progression of the food bolus through the alimentary canal. If CDIAL also inhibits the 5-HT-dependent contractions in the midgut, then larval mosquitoes treated with CDIAL may experience digestive failure. Future studies are required to confirm effects on larval midgut contractions.

At least for CDIAL, we also found a strong inhibition of spontaneous contractions in the hindgut. In larval and adult mosquitoes, the hindgut plays a key role in excretion by reabsorbing solutes and water from the primary urine produced by Malpighian tubules (Piermarini, 2016). Moreover, the visceral muscular contractions of the hindgut play a key role in moving urine and excrement through the ileum and into the rectum before its expulsion. Thus, mosquitoes treated with CDIAL may experience osmoregulatory and/or excretory failure. Additional studies are required to confirm that the inhibition of visceral muscle activity by CDIAL observed *in vitro* in the present study also occurs *in vivo*.

### Hypothesized mode of insecticidal action for CDIAL

The results of the present study suggest that CDIAL inhibits crop contractions by disrupting intracellular Ca^2+^ homeostasis. Notably, the Ca^2+^ ionophore A23187 was the only pharmacological agent to completely inhibit the spontaneous contractions of the crop and change the shape of the crop (i.e., from elliptical to circular) in a similar manner as CDIAL. The change in shape was due to contraction of the distal end of the crop towards the proximal end. We interpret these results as CDIAL and A23187 each inducing an overload of intracellular Ca^2+^ leading to a sustained contractile state without relaxation (i.e., a tetanic paralysis). Although other pharmacological agents completely inhibited the crop (i.e., ivermectin, Gd^3+^, nifedipine, SKF-96365, englerin A), they did not affect crop shape, suggesting induction of a sustained relaxed state (i.e., a flaccid paralysis). Consistent with our interpretations, CDIAL did not affect crop shape in the absence of extracellular Ca^2+^. Moreover, the changes in shape induced by CDIAL were blunted by Gd^3+^, implicating plasma membrane Ca^2+^ channels as likely primary or downstream targets of CDIAL. Given that the frequency of crop contractions was not affected by ryanodine or thapsigargin, and stimulated by chlorantraniliprole, it is unlikely that CDIAL directly agonizes or inhibits intracellular Ca^2+^ transport systems in the crop.

The specific plasma membrane Ca^2+^ channels that are directly or indirectly involved with the apparent tetanic paralysis induced by CDIAL remain to be determined. However, they are unlikely to be of the TRPV or TRPC (a.k.a TRPγ in insects) varieties given pymetrozine’s lack of effects on crop contraction frequency and the apparent flaccid paralysis induced by englerin A, respectively. The complete inhibition of crop contractions by 10 μM Gd^3+^ indicates the likely presence of ORAI and/or voltage-gated Ca^2+^ channels (Putney, 2010), but whether these are affected by CDIAL remains to be determined. Additional molecular, pharmacological, and reverse genetic studies will be required to elucidate the specific biochemical targets involved with the tetanic paralysis induced by CDIAL.

### Insights into the regulation and mechanisms of crop contractile activity

#### Negative regulation of crop contractions via glutamate

Among the neurotransmitters screened, glutamate was the most effective inhibitor of crop contractile activity. In comparison, acetylcholine and GABA had weak inhibitory effects, while histamine and aspartate were without detectable effects. The strong inhibitory effects of glutamate were most likely mediated by GluCl channels given the strong inhibition of crop contractions by ivermectin and ibotenate, which are known agonists of dipteran GluCl channels (Atif et al., 2020; Cully et al., 1996; Eguchi et al., 2006; Fuse et al., 2016; Meyers et al., 2015). When activated under physiological conditions GluCl channels mediate an inward Cl^-^ conductance that hyperpolarizes nerve or muscle cells, thereby reducing the chances of an action potential and muscle activity (Wolstenholme, 2012). Consistent with this mode of action, despite completely inhibiting crop contractions, neither ivermectin nor ibotenate affected the shape of the crops, suggesting a flaccid paralysis wherein the muscle enters a sustained relaxed state. The present study is the first to demonstrate pharmacological evidence for GluCl channels in the crop of a dipteran. Future work will be required to confirm the molecular/biochemical presence of these channels in the mosquito crop.

#### Positive regulation of crop contractions via OA

The strong activation of crop contractile activity by OA and amitraz suggest the presence of OA receptors that stimulate myogenic activity when activated. These results contrast with those previously described in the crop of *D. melanogaster* where OA inhibited crop contractile activity (Solari et al., 2017), suggesting distinct roles of OA in regulating the crops of mosquitoes vs. fruit flies. In other insect systems, OA is a well-characterized modulator of visceral muscle activity. In the oviducts and/or bursa of *Locusta migratoria, Rhodnius prolixus, Stomoxys calcitrans*, and *D. melanogaster*, OA inhibits myogenic activity by causing a rise of intracellular cAMP (Cook and Wagner, 1992; Hana and Lange, 2017a, b; Lange and Orchard, 1986; Lange and Tsang, 1993; Nykamp and Lange, 2000; Orchard and Lange, 1985; Rodríguez-Valentín et al., 2006). On the other hand, in the spermathecae of *L. migratoria*, oviducts of *Gryllus bimaculatus* and *Ae. aegypti*, ovarian peritoneal sheath of *D. melanogaster*, and hindgut of *Ae. aegypti*, OA stimulates myogenic activity via an increase of intracellular Ca^2+^ and/or cAMP (Clark and Lange, 2003; Messer and Brown, 1995; Middleton et al., 2006; Sefiani, 1987; Tamashiro and Yoshino, 2014b). The varying physiological effects of OA among different tissues and species is likely a consequence of differential expression of OA receptor sub-types that signal to particular secondary messengers (Evans and Maqueira, 2005; Evans and Robb, 1993; Farooqui, 2012).

#### Stimulatory effects of 5-HT and OA

Taking the results of the present study together with those in our previous study (Calkins et al., 2017) indicate that the crop of *Ae. aegypti* has at least two physiological pathways for stimulating myogenic activity: serotonergic and octopaminergic. A previous study on the oviduct and hindgut of *Ae. aegypti* also found that 5-HT and OA each elicited stimulatory effects on myogenic activity (Messer and Brown, 1995). Thus, 5-HT and OA may play redundant physiological roles in mosquito visceral muscle. This is not always the case in insects. For example, in the locust oviduct, 5-HT stimulated myogenic activity (Lange, 2004), whereas OA inhibited it (Orchard and Lange, 1985). We have previously shown that the stimulation of myogenic activity in the mosquito crop by 5-HT is likely mediated by an increase of intracellular cAMP (Calkins et al., 2017); whether OA stimulates the crop via a similar or distinct mechanism remains to be determined. At least in the oviduct of *G. bimaculatus*, OA appears to signal to the opening of ryanodine receptors, which when activated increase the frequency of visceral muscle contractions (Tamashiro and Yoshino, 2014a, b). The stimulatory effect of chlorantraniliprole (a ryanodine receptor agonist) on the contraction frequency in the present study suggests that a similar stimulatory pathway may be present in the mosquito crop. Whether OA and/or 5-HT utilize this pathway in the crop and/or other mosquito visceral muscles remains to be determined. It would also be interesting to determine if the agonistic effects of UGAN on crop contractions found in the present study are mediated by influencing a biochemical target in the OA and/or 5-HT stimulatory pathway(s).

### Conclusions

In summary, the present study provides the first insights into the insecticidal mode of action of CDIAL and other drimane sesquiterpene dialdehydes against mosquitoes. CDIAL’s complete inhibition of the spontaneous, 5-HT-stimulated and OA-stimulated contractions of the crop indicate disruption of a mechanism central to visceral muscle function. Collectively our data lead us to generate the hypothesis that CDIAL’s disruption of visceral muscle activity is due to the onset of a tetanic paralysis caused by an overloading of intracellular Ca^2+^ via the activation of plasma membrane Ca^2+^ channels. We also found that CDIAL inhibited the spontaneous contractile activity of the mosquito hindgut suggesting a potentially universal disruption to visceral muscle in the alimentary canal that might lead to digestive, osmoregulatory, and/or excretory failure in mosquitoes *in vivo*. The present study also provides the first pharmacological evidence for putative roles of GluCl channels and OA receptors in regulating myogenic activity of the mosquito crop. The results of our study will guide future research aiming to develop natural product-inspired insecticides with novel modes of action and elucidate the molecular physiology of the mosquito crop.

## Acknowledgments

The authors thank Drs. Ryan Arvidson (College of Wooster), Reed Johnson (The Ohio State University), Angela Lange (University of Toronto Mississagua) and Ian Orchard (University of Toronto Mississauga) for valuable conversations about the present study. The authors are also grateful to Dr. John Beutler (National Cancer Institute, Molecular Targets Program, Bethesda, MD) who generously provided the englerin A used in the present study, and Mr. Erick Martinez Rodriguez and Ms. Nuris Acosta (The Ohio State University) for technical support.

