## Supplemental Tables and Figures for "Stop the crop: insights into the insecticidal mode of action of cinnamodial against mosquitoes"

Table S1. Pharmacological agents utilized in the present study. All were obtained from Thermo Fisher Scientific except for pilocarpine (VWR International, Radnor, PA), carbaryl (Sigma-Aldrich, St. Louis, MO), ibotenic acid (Sigma-Aldrich), propoxur (Sigma-Aldrich), and englerin A (gift of Dr. John Beutler, National Cancer Institute, Molecular Targets Program, Bethesda, MD).

| Chemical name | Abbreviation or name used in text | Solvent | Description | Supporting References |
|---|---|---|---|---|
| **Neurotransmitters** | | | | |
| 4-aminobutyric acid | GABA | $dH_2O$ | Neurotransmitter | |
| 5-hydroxytryptamine | 5-HT | 0.1 N NaOH | Biogenic amine; stimulates contractions in mosquito crops | (Calkins et al., 2017) |
| Acetylcholine chloride | Acetylcholine | DMSO | Neurotransmitter | |
| Amitraz | | DMSO | Octopamine receptor agonist | (Kita et al., 2017) |
| Histamine dihydrochloride | Histamine | $dH_2O$ | Neurotransmitter | |
| L-aspartic acid | Aspartate | 0.1 N NaOH | Neurotransmitter | |
| L-glutamic acid | Glutamate | 0.1 N NaOH | Neurotransmitter | |
| Octopamine hydrochloride | OA | $dH_2O$ | Biogenic amine; inhibits contractions in fly crops | (Solari et al., 2017) |
| **Glutamate receptor modulators** | | | | |
| Ibotenic acid | Ibotenate | 0.1 N NaOH | Glutamate-gated $Cl^-$ channel agonist | (Cully et al., 1996) |
| Ivermectin | Ivermectin | DMSO | Glutamate-gated $Cl^-$ channel agonist | (Atif et al., 2020; Cully et al., 1996; Eguchi et al., 2006; Fuse et al., 2016; Meyers et al., 2015) |

| | | | | |
|---|---|---|---|---|
| Kainic acid | Kainate | 0.1 N NaOH | Ionotropic glutamate receptor agonist | (Han et al., 2015; Li et al., 2016) |
| L-quisqualic acid | Quisqualate | 0.1 N NaOH | Ionotropic glutamate receptor agonist | (Han et al., 2015; Li et al., 2016) |
| Muscimol | Muscimol | $dH_2O$ | Ionotropic glutamate receptor agonist | (Hosie and Sattelle, 1996; McGonigle and Lummis, 2010) |
| N-Methyl-D-aspartic acid | NMDA | $dH_2O$ | Ionotropic glutamate receptor agonist | (Xia et al., 2005) (Bhatt and Cooper, 2005) |
| **Acetylcholine receptor modulators** | | | | |
| Imidacloprid | Imidacloprid | DMSO | Nicotinic acetylcholine receptor agonist | (Zhuang et al., 2016) |
| Pilocarpine | Pilocarpine | DMSO | Muscarinic acetylcholine receptor agonist | (Gross and Bloomquist, 2018) |
| **Acetylcholinesterase inhibitors** | | | | |
| Carbaryl | Carbaryl | DMSO | Acetylcholinesterase inhibitor | (Anderson and Coats, 2012) |
| Carvacrol | Carvacrol | DMSO | Acetylcholinesterase inhibitor | (Anderson and Coats, 2012) |
| Propoxur | Propoxur | DMSO | Acetylcholinesterase inhibitor | (Engdahl et al., 2015; Swale et al., 2015) |
| **$Ca^{2+}$ and TRP channel modulators** | | | | |
| Englerin A | Englerin A | DMSO | TRPC channel agonist | (Akbulut et al., 2015; Carson et al., 2015) |
| $GdCl_3 \cdot 6 H_2O$ | $Gd^{3+}$ | $dH_2O$ | $Ca^{2+}$ channel blocker | (Jörs et al., 2006) |

| | | | | |
|---|---|---|---|---|
| Nifedipine | Nifedipine | DMSO | $Ca^{2+}$ channel blocker | (Browne and O'Donnell, 2018; Dube et al., 2000; Wegener and Nässel, 2000) |
| Pymetrozine | Pymetrozine | DMSO | TRPV channel agonist | (Nesterov et al., 2015) |
| SKF-96365 hydrochloride | SKF-96365 | DMSO | $Ca^{2+}$ channel blocker | (Chen et al., 2013) |
| **Modulators of intracellular $Ca^{2+}$ homeostasis** | | | | |
| A23187 | A23187 | DMSO | $Ca^{2+}$ ionophore | (Dube et al., 2000) |
| Chlorantraniliprole | Chlorantraniliprole | DMSO | Ryanodine receptor agonist | (Cordova et al., 2006; Lahm et al., 2007) |
| Ryanodine | Ryanodine | DMSO | Ryanodine receptor inhibitor | (Wegener and Nässel, 2000) |
| Thapsigargin | Thapsigargin | DMSO | Intracellular $Ca^{2+}$-ATPase inhibitor | (MacPherson et al., 2001; Tamashiro and Yoshino, 2014a; Yu and Beyenbach, 2002) |

### Supplemental Figures

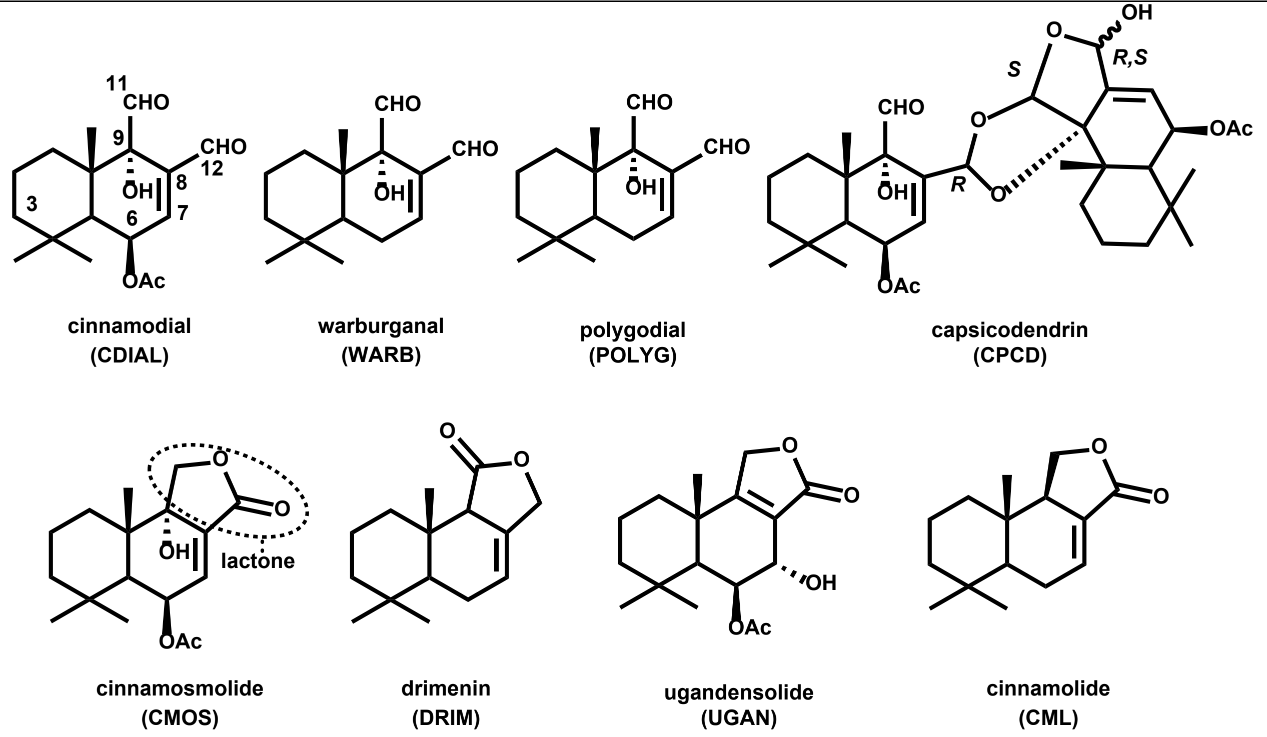


**Supplemental Fig. 1**. Structures of natural drimane sesquiterpenes utilized in the present study. Carbon atoms in CDIAL are numbered. Previous studies by our group have demonstrated that CDIAL, WARB, POLYG, and CPCD possess relatively strong mosquitocidal activity compared to CMOS, DRIM, UGAN, and CML(Inocente et al., 2019; Inocente et al., 2018).

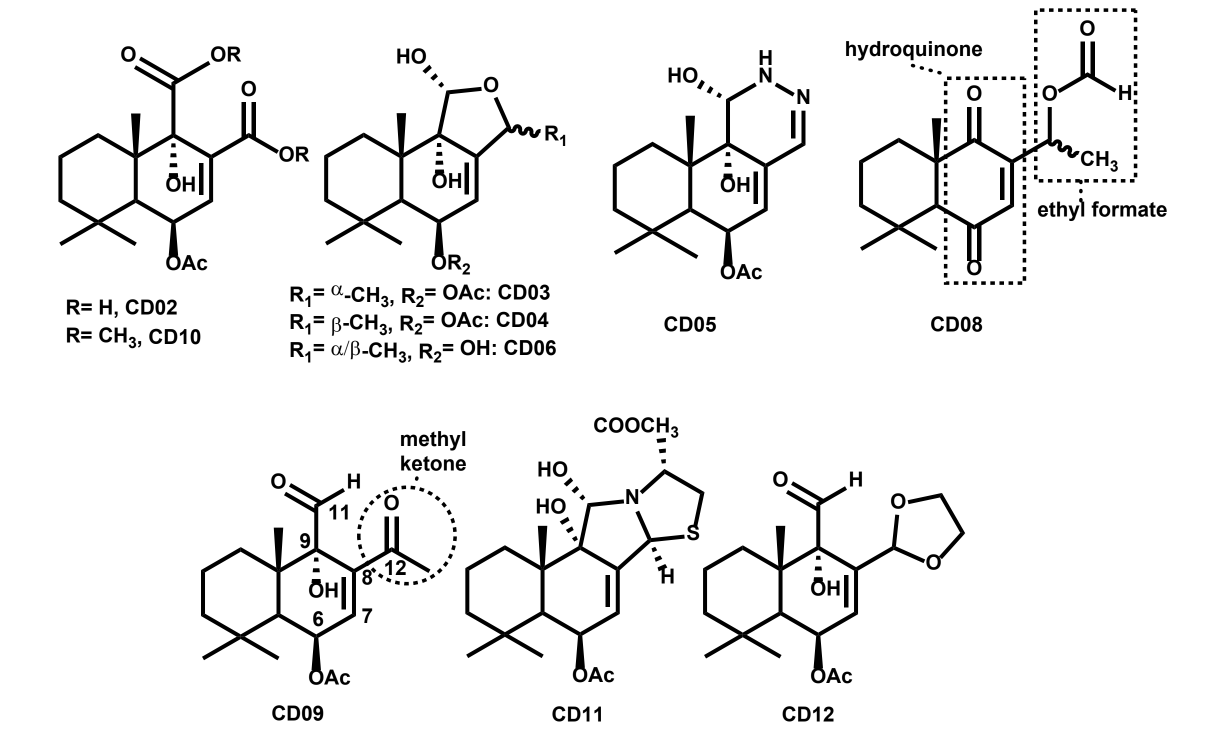


**Supplemental Fig. 2**. Structures of semi-synthetic CDIAL derivatives used in the present study. Carbon atoms are numbered in CD09. We have previously demonstrated that CD08 and CD09 possess similar or more potent mosquitocidal activity than CDIAL (Manwill et al., 2020).

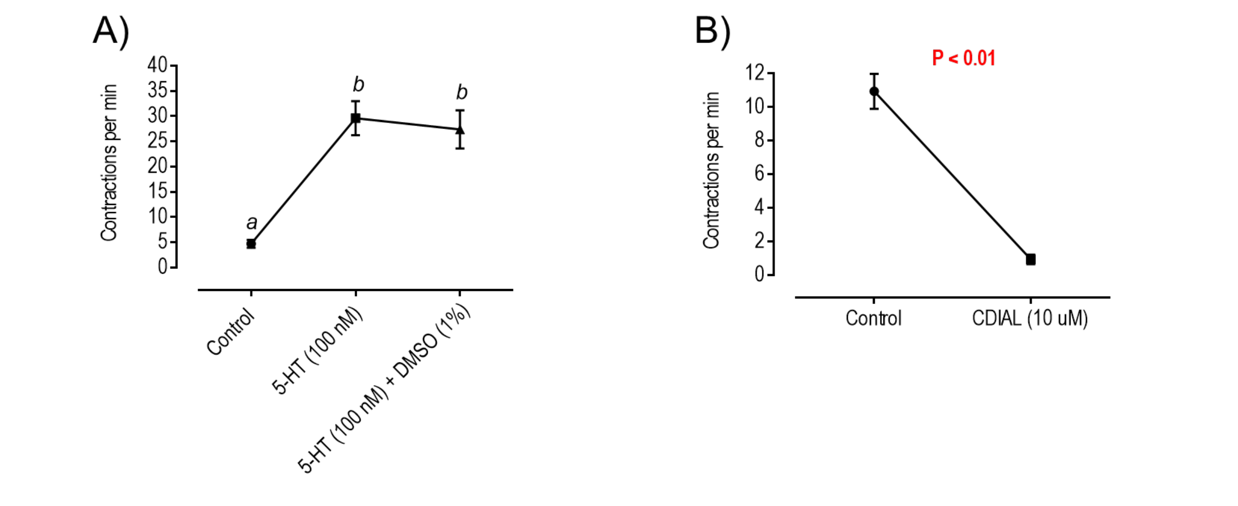


Supplemental Fig.3. A) Effects of 1% DMSO (CDIAL solvent) *in vitro* on the rate of crop contractions stimulated by 5-HT (100 nM). Values are means ± SEM; N = 5. Lower-case letters indicate statistical categorization of the means as determined by a repeated measures one-way ANOVA and Tukey's multiple comparisons test ($P < 0.05$). B) Effects of CDIAL *in vitro* on spontaneous contractions of the hindgut. Values are means ± SEM; N = 5. *P* value from a paired t-test is indicated.

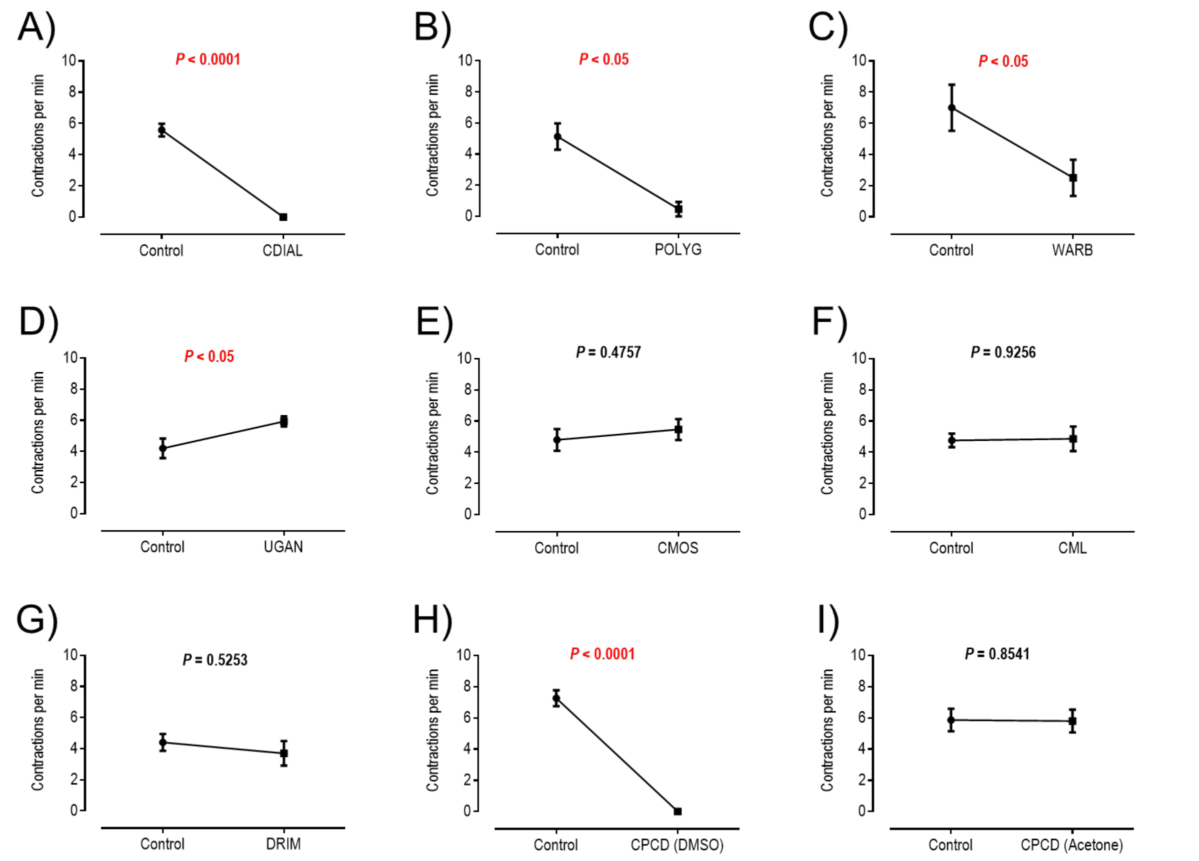


Supplemental Fig.4. Effects of natural CDIAL derivatives *in vitro* on spontaneous contraction rates of the crop. All derivatives were tested at 10 µM and the final solvent concentration was 1% DMSO, except for 'CPCD (acetone)' where the final solvent concentration was 1% acetone. Values are means ± SEM. *P* values from paired t-tests are indicated. N = 19 (CDIAL), 5 (POLYG), 6 (WARB), 5 (UGAN), 10 (CMOS), 10 (CML), 10 (DRIM), 5 (CPCD DMSO), and 5 (CPCD Acetone).

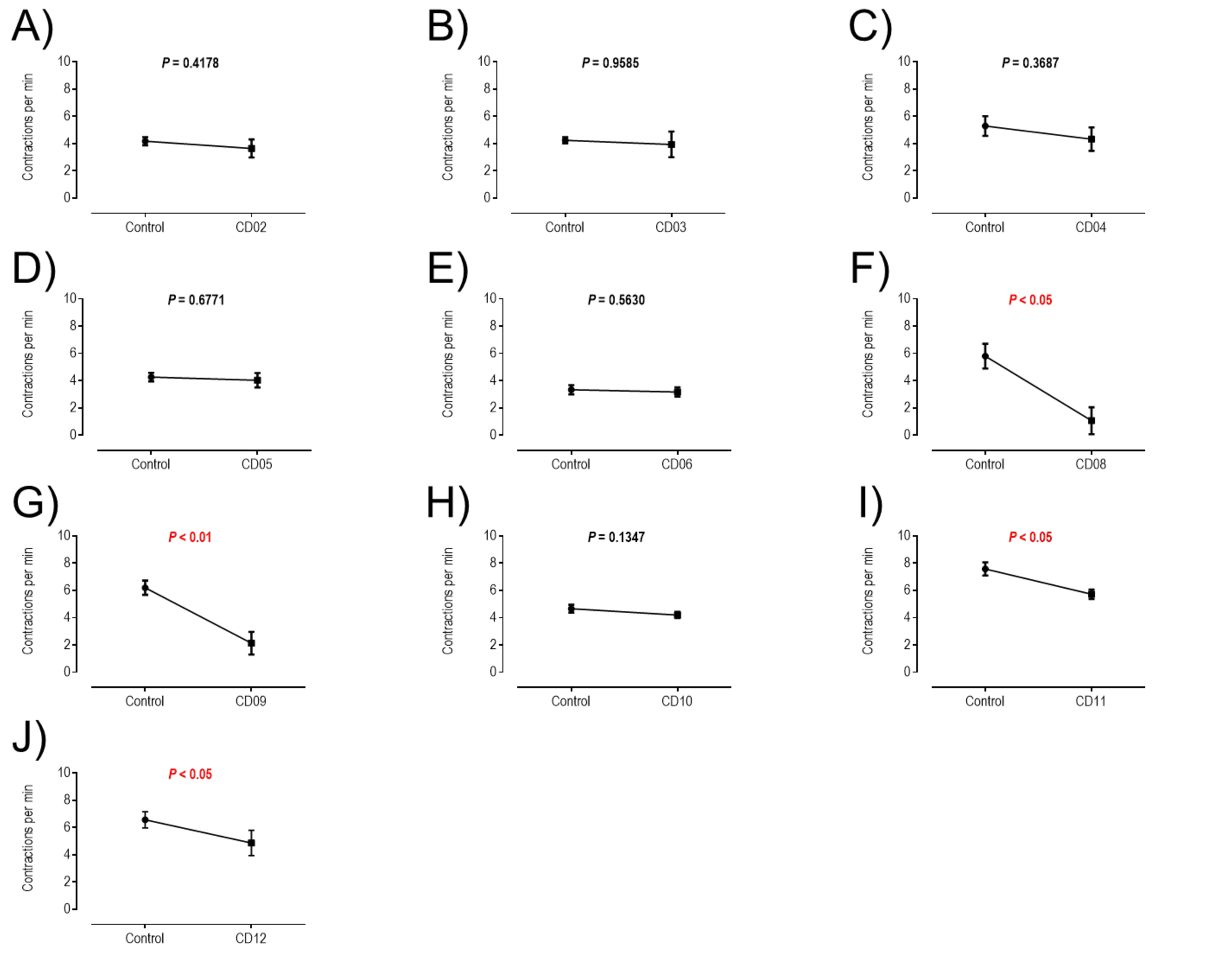


Supplemental Fig.5. Effects of semi-synthetic CDIAL derivatives *in vitro* on spontaneous contraction rates of the crop. All derivatives were tested at 10 µM and the final solvent concentration was 1% DMSO. Values are means ± SEM. *P* values from paired t-tests are indicated. N = 18 (CD02), 12 (CD03), 9 (CD04), 10 (CD05), 8 (CD06), 5 (CD08), 10 (CD09), 5 (CD10), 12 (CD11), and 10 (CD12).

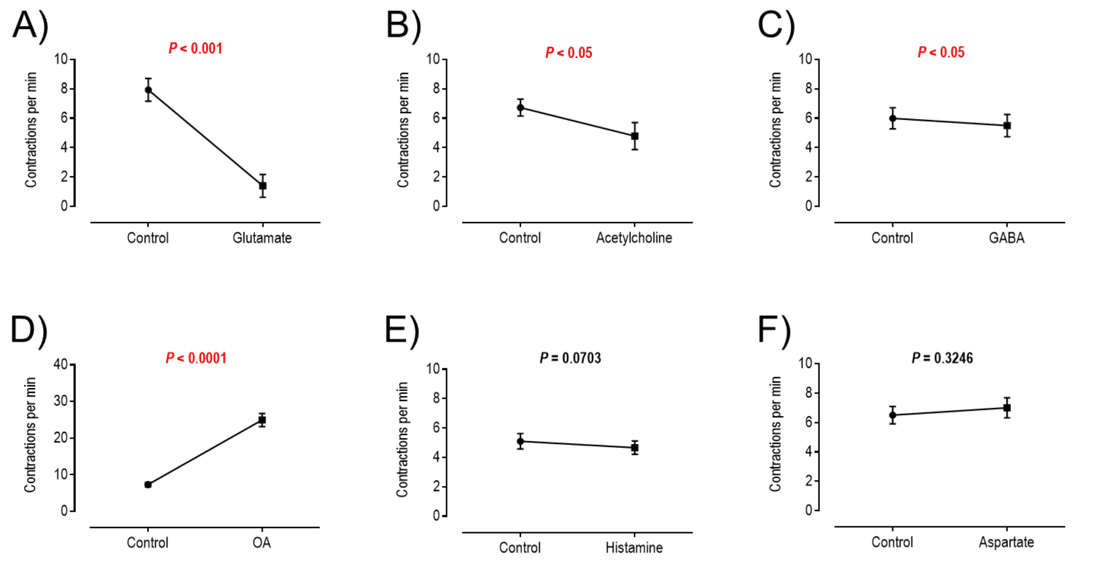


Supplemental Fig.6. Effects of neurotransmitters *in vitro* on spontaneous contraction rates of the crop. All were tested at 1 mM and the final solvent concentration was 1% of the corresponding solvent in Supplemental Table 1. Values are means ± SEM. *P* values from paired t-tests are indicated. N = 10 for each compound.

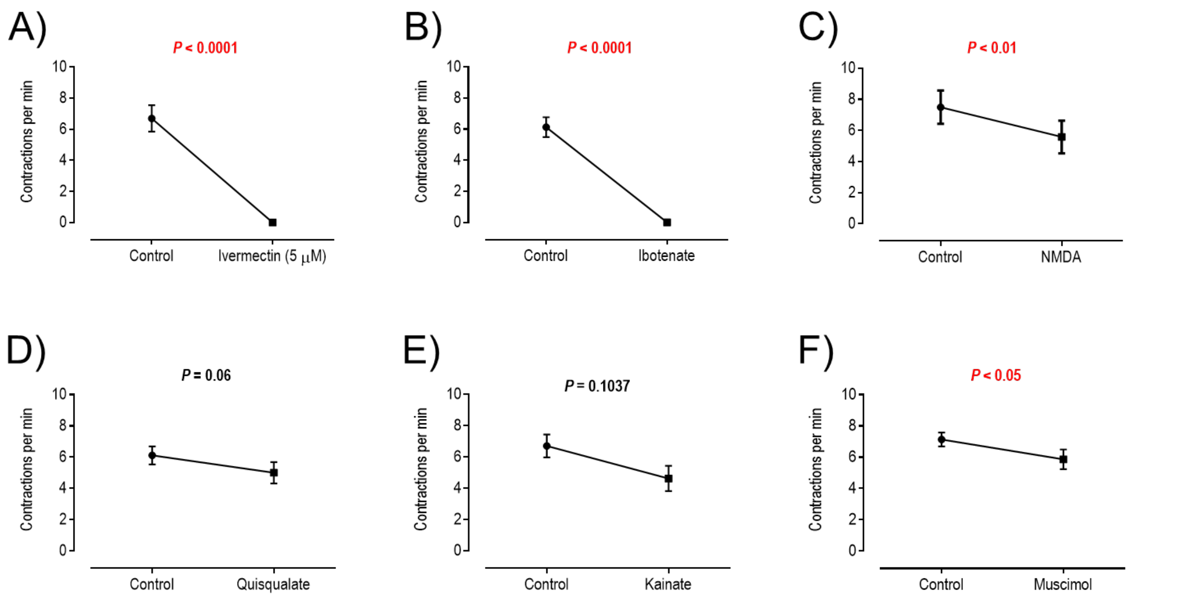


Supplemental Fig.7. Effects of agonists of glutamate-gated chloride channels (A-B), ionotropic glutamate receptors (C-E), or GABA receptors (F) *in vitro* on spontaneous contraction rates of the crop. All were tested at 1 mM, except for ivermectin (5 μM). The final solvent concentration was 1% of the corresponding solvent in Supplemental Table 1. Values are means ± SEM. *P* values from paired t-tests are indicated. N = 10 for ivermectin, ibotenate, and muscimol. N = 8 for NMDA and kainite. N = 9 for quisqualate.

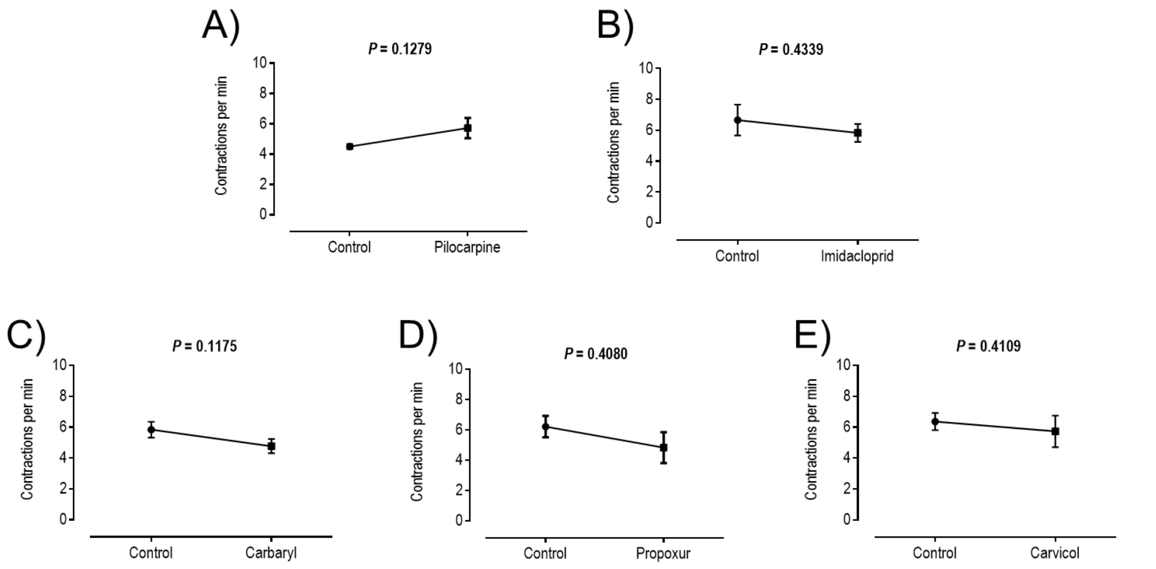


Supplemental Fig.8. Effects of agonists of acetylcholine receptors (A-B) or inhibitors of acetylcholinesterase (C-E) *in vitro* on spontaneous contraction rates of the crop. All were tested at 10 μM, except for carbaryl (100 μM). The final solvent concentration was 1% of the corresponding solvent in Supplemental Table 1. Values are means ± SEM. *P* values from paired t-tests are indicated. N = 10 for all compounds except pilocarpine and propoxur where N = 6.

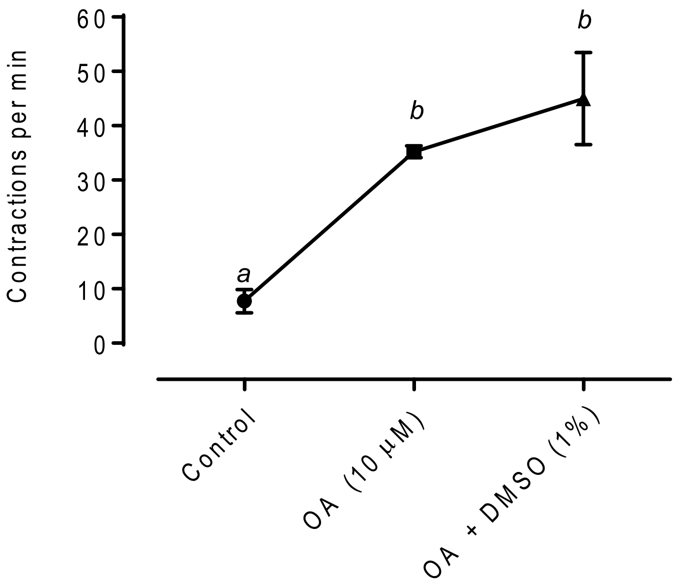


Supplemental Fig.9. Effects of 1% DMSO (CDIAL solvent) *in vitro* on the rate of crop contractions stimulated by OA (10 µM). Values are means ± SEM; N = 5. Lower-case letters indicate statistical categorization of the means as determined by a repeated measures one-way ANOVA and Tukey's multiple comparisons test ($P < 0.05$).

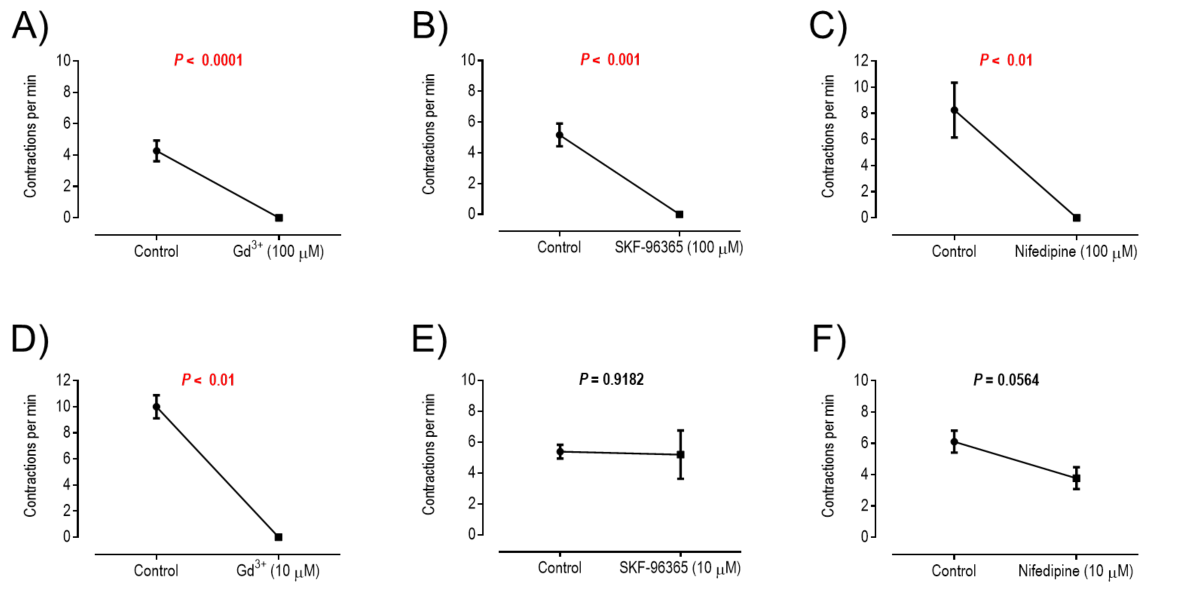


Supplemental Fig.10. Effects of blockers of plasma membrane $Ca^{2+}$ channels *in vitro* on spontaneous contraction rates of the crop. Compounds were tested at 100 µM (A-C) or 10 µM (D-F). The final solvent concentration was 1% of the corresponding solvent in Supplemental Table 1. Values are means ± SEM. *P* values from paired t-tests are indicated. N = 11, 4, or 6 for 100 µM $Gd^{3+}$, SKF-96365, or nifedipine, respectively. N = 3, 7, or 9 for 10 µM $Gd^{3+}$, SKF-96365, or nifedipine, respectively.

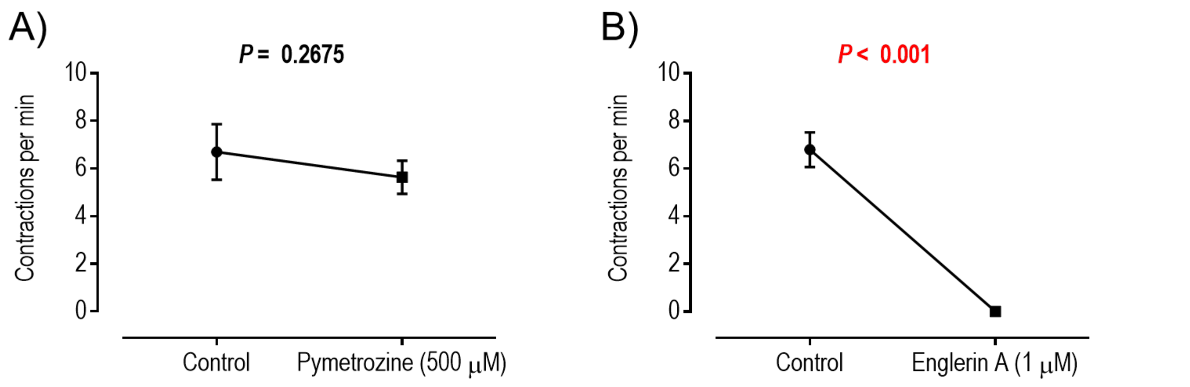


Supplemental Fig.11. Effects of agonists of TRPV (A) or TRPC (B) channels *in vitro* on spontaneous contraction rates of the crop. The final solvent concentration was 1% of the corresponding solvent in Supplemental Table 1. Values are means ± SEM. *P* values from paired t-tests are indicated. N = 10 for pymetrozine; N =5 for englerin A.

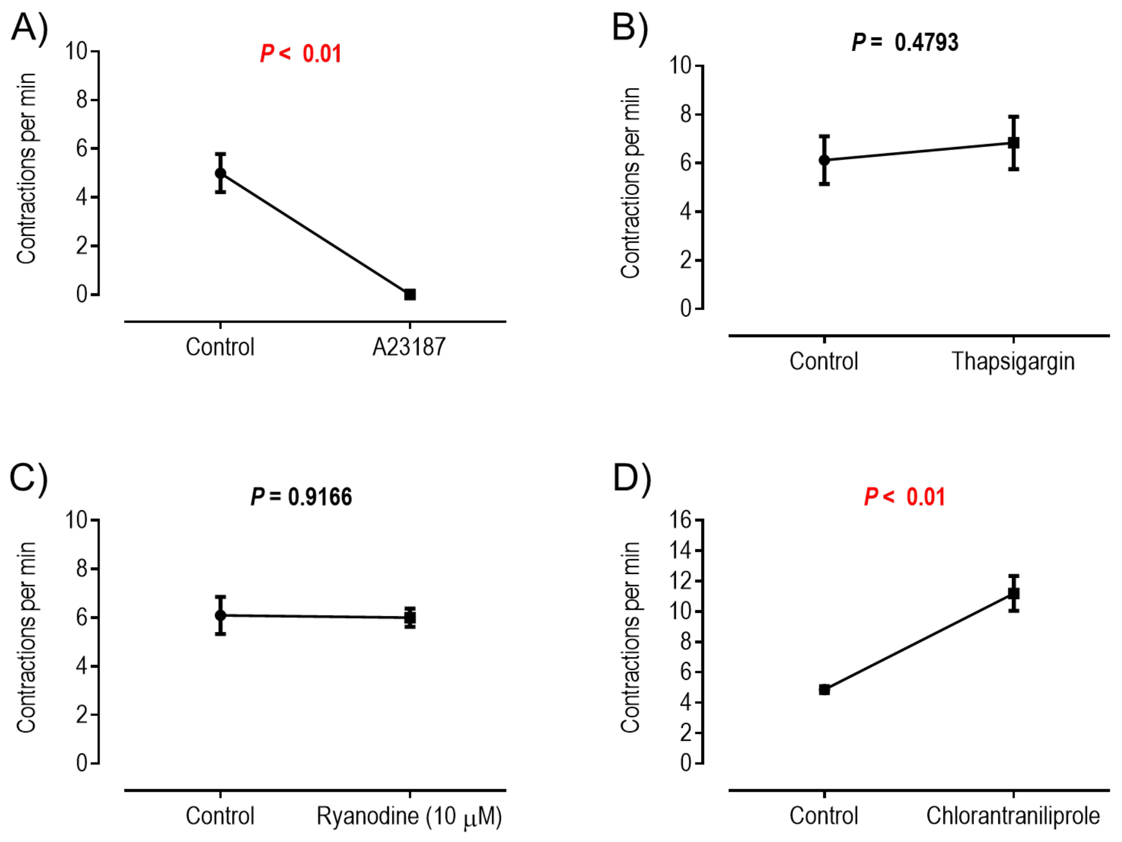


Supplemental Fig.12. Effects of modulators of intracellular $Ca^{2+}$ homeostasis *in vitro* on spontaneous contraction rates of the crop. All were tested at 1 μM except for ryanodine (10 μM). The final solvent concentration was 1% of the corresponding solvent in Supplemental Table 1. Values are means ± SEM. *P* values from paired t-tests are indicated. N = 5 for A23187 and chlorantraniliprole. N = 8 for thapsigargin and N = 7 for ryanodine.

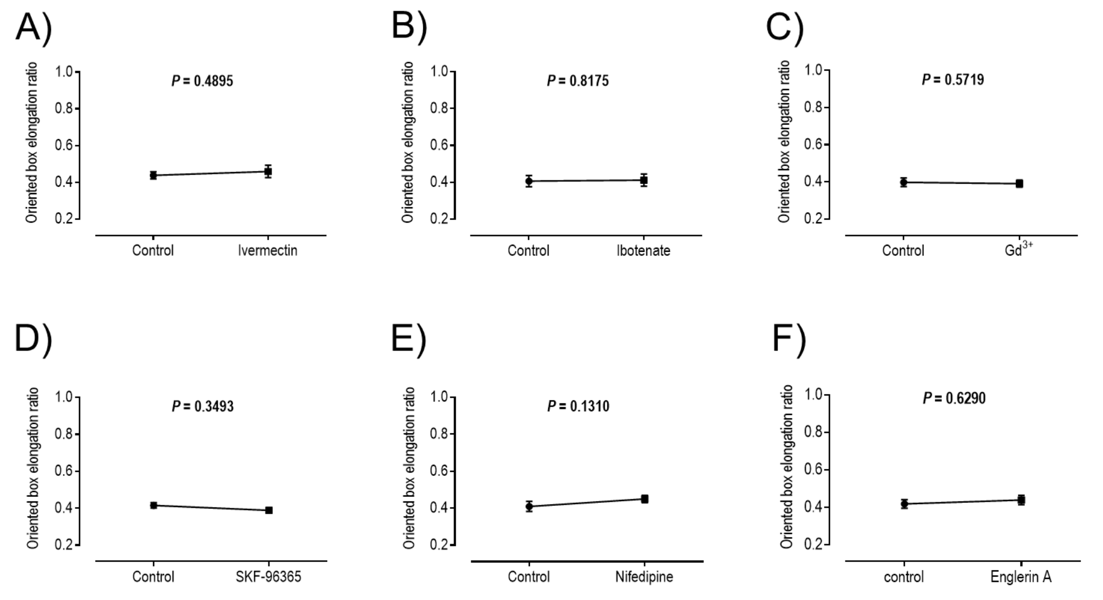


Supplemental Fig.13. Effects of GluCl agonists (A-C), $Ca^{2+}$ channel blockers (C-E), or a TRPC channel agonist (F) on the OBER of crops. Values are means ± SEM. P values from paired t-tests are indicated. N = 5 for each compound.
